# The evolution of germ-soma specialization under different genetic and environmental effects

**DOI:** 10.1101/2021.04.02.438224

**Authors:** Denis Tverskoi, Sergey Gavrilets

## Abstract

Division of labor exists at different levels of biological organization - from cell colonies to human societies. One of the simplest examples of the division of labor in multicellular organisms is germ-soma specialization, which plays a key role in the evolution of organismal complexity. Here we formulate and study a general mathematical model exploring the emergence of germ-soma specialization in colonies of cells. We consider a finite population of colonies competing for resources. Colonies are of the same size and are composed by asexually reproducing haploid cells. Each cell can contribute to activity and fecundity of the colony, these contributions are traded-off. We assume that all cells within a colony are genetically identical but gene expression is affected by variation in the microenvironment experienced by individual cells. Through analytical theory and evolutionary agent-based modeling we show that the shape of the trade-off relation between somatic and reproductive functions, the type and extent of variation in within-colony microenvironment, and, in some cases, the number of genes involved, are important predictors of the extent of germ-soma specialization. Specifically, increasing convexity of the trade-off relation, the number of different environmental gradients acting within a colony, and the number of genes (in the case of random microenvironmental effects) promote the emergence of germ-soma specialization. Overall our results contribute towards a better understanding of the role of genetic, environmental, and microenvironmental factors in the evolution of germ-soma specialization.

## 1 Introduction

The history of life on Earth can be viewed as a sequence of major evolutionary transitions to more complex hierarchically nested levels of biological organization (Smith and Szathmary 1995; Carroll 2001; Grosberg and Strathmann 2007) including transitions from self-catalyzed chemical reactions to the RNA-world, from genes to prokaryotic cells, from prokaryotic to eukaryotic cells, from single cells to multicellular organisms, and from discrete multicellular individuals to social groups. These transitions were accompanied by the emergence of division of labor between lower-level units which contribute to fitness of the new higher-level individual (Michod 1996, 1997, 2007).

Division of labor is found at different levels of biological organization. Examples include specialization of duplicated genes (Reuffler and Wagner 2012), germ-soma separation in Volvocales (Kirk 2003; Nedelcu and Michod 2006), cyanobacteria (Rossetti *et al*. 2010) and hydrozoans (Rossetti *et al*. 2010), specialization on various non-overlapping metabolic processes like carbon fixation and nitrogen fixation in cyanobacteria (Fay 1992; Berman-Frank *et al*. 2003; Flores and Herrero 2010), emergence of casts in social insects (Robinson 1992; Beshers and Fewell 2001; Wilson 2008), sexual division of labor in small hunter-gatherer societies (Panter-Brick 2002), and division of labor within complex organizational entities such as firms and markets (Smith 2008; Rodriguez-Clare 1996; Stigler 1951).

Division of labor between reproductive and non-reproductive functions, also known as germ-soma specialization (Folse and Roughgarden 2010), is the earliest observable example of labor division within multicellular groups. Germ-soma specialization leads to the emergence of reproductive altruism, a phenomenon, where a fraction of cells, called somatic, contribute to the colony’s functionality but cannot produce an offspring themselves (Nedelcu 2009; Goldsby *et al.* 2014). Germ-soma specialization (and reproductive altruism as well) plays a key role in promoting the evolution of organismic complexity (Yanni *et al.* 2020), limiting reversion to unicellularity (Libby and Ratcliff 2014), or even facilitating the emergence of eusociality (Nedelcu 2009), However its origin, mechanics and evolution remain understudied (McShea 2000; Yanni *et al.* 2020).

A number of empirical studies have shown that environmental conditions can impact the level of germ-soma specialization. For example, Bonner (2003) explored the effect of environmental conditions on the level of germ-soma specialization in slime molds. In case of resources scarcity, the unicellular individuals assemble together and combine into multicellular aggregates, where they split into the somatic stalk and reproductive spores. Numbers of somatic and reproductive cells vary in accordance with changes in temperature, oxygen concentration, or other chemical gradients. Nedelcu (2009) showed that various environmental factors such as limited nutrient concentration, water stress or cold can cause reproductive suppression in Volvocales. She argued that a temporally varying environment promotes the evolution of reproductive altruism in Volvocales. Herron *et al*. (2014) showed that the type of environment (still or mixed) has a statistically significant effect on the proportion of somatic cells in *Pleodorina starrii* colonies.

Because the underlying biological processes are very complex, mathematical modeling can deepen our understanding of the evolution of germ-soma specialization. Earlier modeling work has focused on the effects of trade-offs between reproductive and somatic functions (Michod *et al*. 2006; Leslie *et al.* 2017), group size (Michod *et al.* 2006), genetic relatedness (Cooper and West 2018), developmental plasticity (Gavrilets 2010), positional effects (Reuffler and Wagner 2012) and topological constraints (Yanni *et al.* 2020) on the dynamics of labor division. Some researchers studied how environmental factors influence the emergence of germ-soma specialization. In particular, Willensdorfer (2009) assumed that cells contribute to the acquisition of a limited resource that is needed by reproductive cells for growth. He showed that in such a situation complete germ-soma specialization emerges in small colonies with a small number of reproductive cells. Rossetti *et al*. (2010) presented a detailed model for studying the emergence of differentiate multicellularity in cyanobacteria, where it was assumed that colonies compete with each other for solar energy, which is subsequently converted into chemical energy (fixed carbon). Tverskoi *et al.* (2018) focused on the impact of environmental factors, positional effects, and the trade-off curvature on germ-soma specialization in a cell colony. Since the model introduced by Tverskoi *et al.* (2018) is static, it tells us little about the dynamics of germ-soma separation, forces that shape this process, and underlying mechanisms.

Here we extend the work of Gavrilets (2010) and Tverskoi *et al.* (2018) by examining effects of environmental factors, positional effects and the trade-off between cell activity and fecundity on the evolution of germ-soma specialization in cell colonies. We posit that the environment affects cell colonies at a between-colony level (via resource-based competition among colonies), and at a within-colony level (via different microenvironmental effects on gene expressions caused by positional effects). We consider the evolution of germ-soma specialization in the case of random microenvironmental effects and in the case of microenvironmental gradients, e.g. due to some chemical gradients within colonies. We seek to find the conditions favouring the evolution of the division of labor and the emergence of a new level of biological organization - a multicellular organism with complete germ-soma differentiation.

## 2 The model

We consider a finite population of colonies each composed by *S* asexually reproducing haploid cells. Time is discrete. Gene expression is affected by variation in microenvironment experienced by individual cells within the colony. Colonies compete for resources. Colonies surviving to the stage of reproduction disintegrate and the released cells start new daughter-colonies which are produced by binary division. Mutation occurs during cell divisions.

### Cell genotype and phenotype

Colonies differ in *G* genes. All cells within a colony are assumed to be genetically identical. The cell’s genotype is specified by a vector *g* with elements *g*_*i*_ scaled to be between 0 and 1 (*i* = 1, …, *G*). These genes control a cell’s activity *a* and fecundity *b* (Michod *et al.* 2006; Gavrilets 2010). The former is a measure of the cell’s contribution towards the colony’s ability to secure resources it requires, e.g. via flagellar action (Kirk 2001; Goldstein 2015; Brumley et al. 2015). The latter is defined as the probability that the cell successfully starts a new colony. Both a and b are restricted to be between 0 and 1.

We define the *i*-th gene expression level *x*_*i*_ = *e*_*i*_*g*_*i*_, where parameter *e*_*i*_ specifies microenvironmental effects (e.g. via effects of some chemical gradients) on gene expression. Microenvironment effects *e*_*i*_ may differ between different cells of the same colony. That is, we use a linear reaction norm for gene expression with zero intercept (de Jong 1990; Gavrilets and Scheiner 1993). Let the cumulative gene effect be *x* = ∑ *e*_*i*_*g*_*i*_ and let the cell’s fecundity be an S-shaped function of *x*, which changes between 0 and 1:

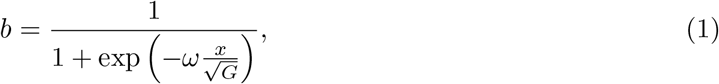

where the division by 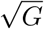 inside the exponential is a technical assumption to remove the dependence of the variation of *b* on the number of genes, and the positive parameter *ω* specifies the steepness of this function.

Following Michod *et al.* (2006), we assume that fecundity and activity within each cell are traded off and write cell activity as *a* = *ϕ*(*b*), where function *ϕ* is continuous and monotonically decreasing on [0, 1] with *ϕ*(0) = 1 and *ϕ*(1) = 0. (See Tverskoi *et al.* (2018) for additional technical requirements on *ϕ*). Our results below will be based on the following form of *ϕ*:

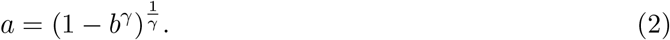

(Egas *et al.* 2004; Leslie *et al.* 2017; de Oliveira *et al.* 2020). Parameter *γ* > 0 controls the shape of the trade-off relation which is convex for *γ <* 1, linear for *γ* = 1, and concave for *γ* > 1. According to Michod (2005, 2006); Michod *et al.* (2006), in volvocine green algae the trade-off function between activity and fecundity is convex-like for large-size colonies, and concave for small-size colonies. Below we will make this assumption when discussing the effects of the colony size on the model dynamics.

### Cell microenvironment and cell prototypes

We assume that cells can differ in certain characteristics, such as size or position in the colony, which can affect their gene expression. For example, in Volvox the *regA* gene acts in small cells (proto-soma) suppressing their reproductive function while the *lag* gene acts in large cells (proto-germ) suppressing motility function (Kirk 2005). In *Pleodorina starrii*, cells are divided into tiers arranged from the anterior to the posterior of a colony that is determined according to the motility direction (Herron *et al.* 2014). We will focus on microenvironmental effects. Specifically, we assume that each colony has groups of cells experiencing the same macro-environment described bu vector *e* = (*e*_1_, …, *e*_*G*_) of microenvironmental effects, cells belonging to different groups are characterized by different vectors *e*. We will say that cells belonging to the same group have the same *prototype*, and cells of different groups have different prototypes. For example, all cells in the internal layer of the colony can experience the same microenvironment (and thus are of the same prototype) and all cells in the external layer of the colony can experience another type of microenvironment (and thus are of a different prototype).

We assume that in the colony there are *s* different prototypes each with the same number of cells. We will consider two ways of specifying microenvironmental effects *e*_*mi*_ (*m* = 1, .., *s*; *i* = 1, .., *G*). In the case of *random microenvironmental effects*, each *e*_*m,i*_ value is drawn randomly and independently from a uniform distribution on [−1, 1]. In the case of *microenvironmental gradients, e*_*m,i*_ values change in a systematic way inside the colony. For example, one can think of a gradient of some chemical along the anterior-posterior axis. In the model, we postulate that |*e*_*m,i*_| decreases geometrically at rate *r* with the distance *m* from the anterior layer, so that *e*_*m*+1,*i*_ = *re*_*m,i*_ with |*e*_1,*i*_| = *d* where *d* > 0 is a constant. We will allow for several different gradients along the same anterior-posterior axes.

### Colony survival

At each time step, colonies compete for a resource of a value *C* > 0. We define the *competitiveness* of the colony as the total activity of its cells: *A* = ∑ _*j*_ *a*_*j*_ and say that the amount of the resource it secures in competition with other colonies is

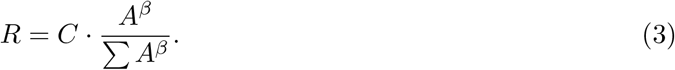

Here the sum in the denominator is over all *N* colonies in the system and a parameter *β* ≥ 1 measures the strength of competition. In the economics literature, this model is known as the Tullock contest (Tullock 1980; Rusch and Gavrilets 2017) and parameter *β* as its decisiveness (Konrad 2009). In volvocine algae, competition in a still environment is stronger than in a mixed environment(Herron *et al.* 2014). This feature can be captured by varying parameter *β*.

We postulate that the colony always survives to the stage of reproduction if the amount of the resource it has secured *R* is higher than a certain threshold *R*_0_. If *R < R*_0_, the probability of survival is

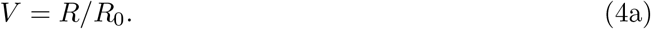

We write the threshold value as

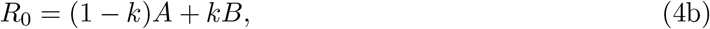

where *B* = ∑*b*_*i*_ is the total fertility. The value of *R*_0_ can be thought of as the total amount of the resource needed to maintain the colony. Parameter 0 *< k <* 1 measures the relative cost of fertility. Thus, in our model the more resources *R* are secured, the more likely the colony survives, and colonies with larger competitiveness *A* and total fertility *B* require more resources. [This model is a stochastic version of a general resource constraint proposed in Tverskoi *et al.* (2018)]. Our formulation implies that the population of colonies is under density-dependent viability selection.

### Reproduction

Each surviving colony disintegrates into *S* independent cells and each cell seeds a new colony of genetically identical cells with the corresponding probability *b*_*i*_. Mutations in an offspring colony genotype *g* happen with probability *µ* per gene. Each mutation modifies the corresponding element of vector *g* by a value drawn randomly and independently from a truncated normal distribution with zero mean and standard deviation *σ*.

### Numerical implementations

To study our model, we used agent-based simulations. The following is a list of parameters which we varied systematically: the number of cells per colony *S* = 8, 16, 32; the number of genes *G* = 2, 8, 32, 128; the strength of competition among colonies *β* = 1, 3, 6; the relative cost of fertility: *k* = 0.1, 0.2, …, 0.9, the shape of the trade-off relation *γ* = 2 for small colonies with size *S* = 8, 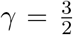 for small colonies with size *S* = 16, and 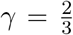 for large colonies with size *S* = 32. The number of gradients in the cell microenvironment *E* was either 1 or 4. We posited that *d* = 10. With *E* = 1, we considered *r* = 0.2, 0.4, 0.6, 0.8. With *E* = 4, we assumed that *r*_1_ = 0.2, *r*_2_ = 0.4, *r*_3_ = 0.6, *r*_4_ = 0.8. We assumed that there are *G/E* genes associated with each gradient. We also assumed that *e*_1,*i*_ = *d* for the first *G/*(2*E*) among the above *G/E* genes, and *e*_*i,i*_ = −*d* for the last *G/*(2*E*) among the above *G/E* genes.

Other parameters used: the number of prototypes *s* = 4; the amount of the resource *C* = 1000; the steepness of an S-shaped function that transforms the cumulative gene effect into cell fecundity *ω* = 4; per gene mutation rate *µ* = 0.001 and per gene mutation standard deviation *σ* = 0.05. Each simulation is run for 10^6^ generations and took averages over the last quarter of generations.

### Cell and colony phenotypes

In our analytical derivations, we defined *a cell phenotype* as its fecundity *b*. Naturally, cells of the same prototype have the same phenotype. However, cells of different prototypes can have the same phenotype as well. Therefore, the number of different cell phenotypes *M* is less than or equal to the number of different prototypes *s* in the colony. The set of all cell phenotypes within the colony is called *a colony phenotype*.

In our simulations, we applied a modification of the single linkage clustering method (Kaufman and Rousseeuw 1990) to the observed distribution of cell fecundities *b* in order to determine the number of different cell phenotypes *M* in the colony. Specifically, we first found the minimum *b*^*min*^ and the maximum *b*^*max*^ values of *b* during the last quarter of time steps for each cell prototype. Then, we posited that two prototypes have the same cell type if the corresponding intervals [*b*^*min*^, *b*^*max*^] intersect or are separated from each other by less than 0.05. This means that cells have the same phenotype, if they have close fecundity values.

### Somatic and reproductive cells

In our simulations, a cell was identified as somatic if *b* ≤ 0.1 (i.e., its contribution to the reproductive task is small). A cell was identified as reproductive if *a* ≤ 0.1 (i.e., its contribution to activity is small). All other cells were classified as unspecialized.

## 3 Results

We studied our model using a combination of analytical approximations focusing on the case of random microenvironmental effects and a very large number of genes, and evolutionary agent-based simulations investigating cases with an arbitrary number of genes *G*. Below we will first present results on random microenvironmental effects. After that we will discuss results on microenvironmental gradients. We will focus on two cases: with a single gradient, and with several gradients.

### 3.1 Random microenvironmental effects

With random microenvironmental effects, the population evolves to an equilibrium at which between-colony variation is very low. Therefore we will focus on the effects of the number of genes *G*, the strength of competition between colonies *β*, and the relative cost of fertility *k* on average characteristics of colonies. Specifically, we will look at the number of different cell phenotypes *M* per colony, the total share *p* of specialized (both reproductive and somatic) cells, the share *p*_*g*_ of specialized reproductive cells, the competitiveness *A*, the total fertility *B*, and the number of surviving colonies *N*.

Our results show that the number of genes *G* and the colony size *S* are key factor controlling the dynamics of the model. However, some general patterns appear regardless of *G* and *S*. First, our numerical simulations show that, in general, the competitiveness *A* of each colony is higher than its total fecundity *B*, except for the cases with small *β* and *k*. Increasing *k* or *β* reduces the total fecundity of a colony and increases its competitiveness (Figure 1a).

**Figure 1:**
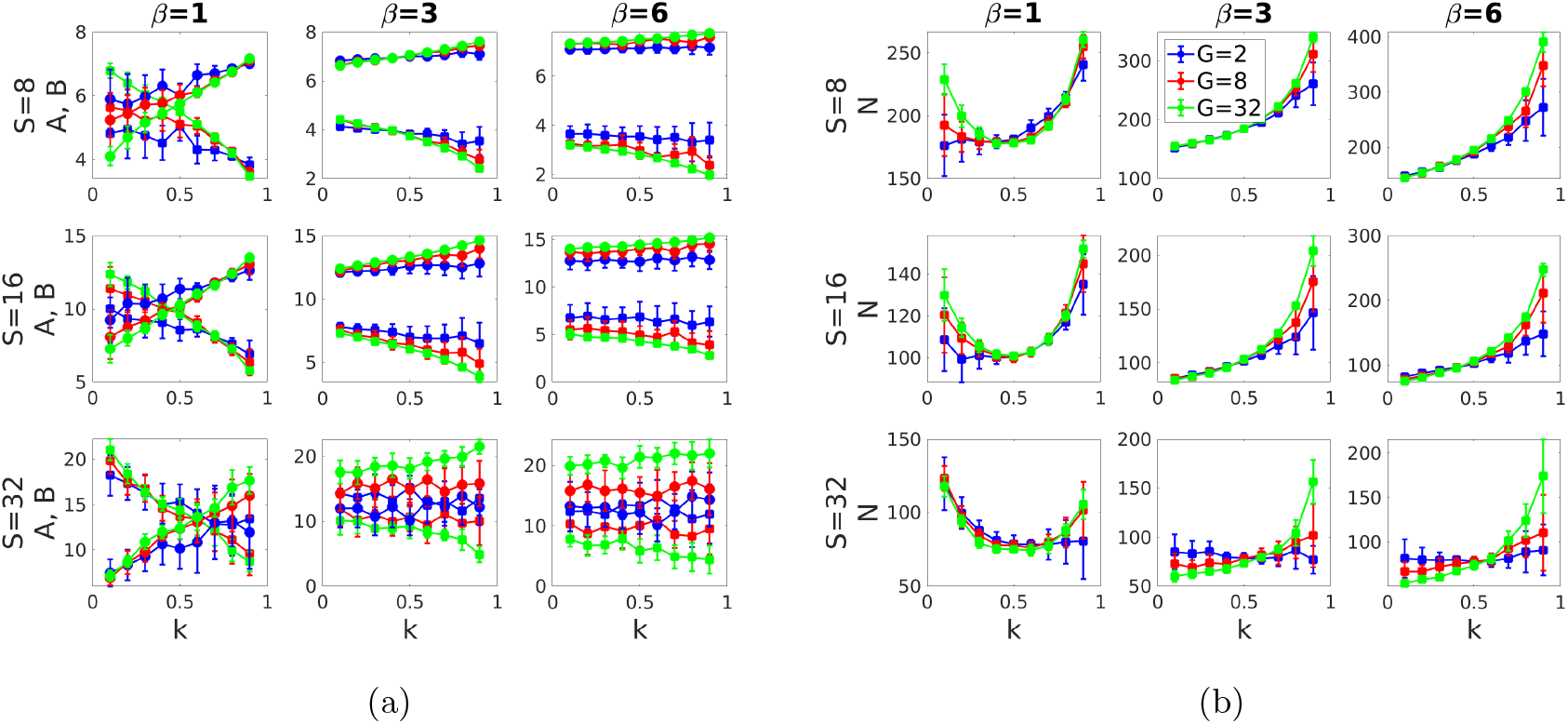
Effects of the relative cost of fecundity *k*, the strength of competition *β*, the number of cells *S* and the number of genes *G* on (a) the competitiveness *A* and the total fecundity *B* of a colony, and (b) the population size *N* in the case of random microenvironmental effects. Baseline parameters: *s* = 4, *C* = 1000, *ω* = 4, *µ* = 0.001 and *σ* = 0.05. Parameter *γ* = 2 for *S* = 8, *γ* = 3*/*2 for *S* = 16 and *γ* = 2*/*3 for *S* = 32. Shown are averages and confidence intervals based on 20 runs each of length 10^6^ time steps for each parameter combination. Results on each run have been calculated based on the last quarter of time steps.

Second, we find that the effect of the strength of competition between colonies *β* on the number of surviving colonies *N* depends on *k*. Increasing *β*: (1) decreases *N*, if *k* is small; (2) has no significant effect on *N*, if *k* is intermediate; and (3) increases *N*, if *k* is large. The effect of the relative cost of fecundity *k* on *N* depends on *β* as well. For small *β* the population size *N* exhibits a U-shaped dependence on *k*. For intermediate or large *β* increasing *k* leads to an increase in *N* . Overall, this implies that large number of surviving colonies are observed in two cases. In the first case, *k* and *β* are small so that colonies contribute more to fecundity than to activity. This results in the large number of offspring colonies. Since *β* is small, the competition between colonies is weak and therefore large number of offspring colonies survive (i.e., *N* is large). In the second case, *k* is large so colonies contribute more to activity than to fecundity. As a result, offspring are more likely to survive during the resource competition (i.e., *N* is large as well). This phenomenon is quite more marked, if the number of genes *G* is large. For more details see Figure 1b.

To explore effects of the number of genes *G* and the colony size *S* we will consider two cases separately: a small number of genes (*G* = 2, 8), and a large number of genes (*G* = 32). First, we will provide results on small-size colonies, and then - on large-size colonies.

#### Small colonies

Using Michod (2005, 2006); Michod *et al.* (2006) conclusions on volvocine green algae we assume that in small colonies the trade-off function describing the relationship between activity *a* and fecundity *b* is concave with *γ* > 1. We will assume that this is the case for colonies with *S* = 8, 16 cells.

If the number of genes is large (i.e. *G* = 32), the population evolves to a state at which all cells have the same unspecialized phenotype (i.e., a colony is characterized by a zero share of specialized cells, *p* = 0). For details see Figure 2c and Figure S1 in the SM. Analytical approximations (for *G* → ∞) show that equilibrium cell fecundity *b*^*\**^ satisfies the following equation:

**Figure 2:**
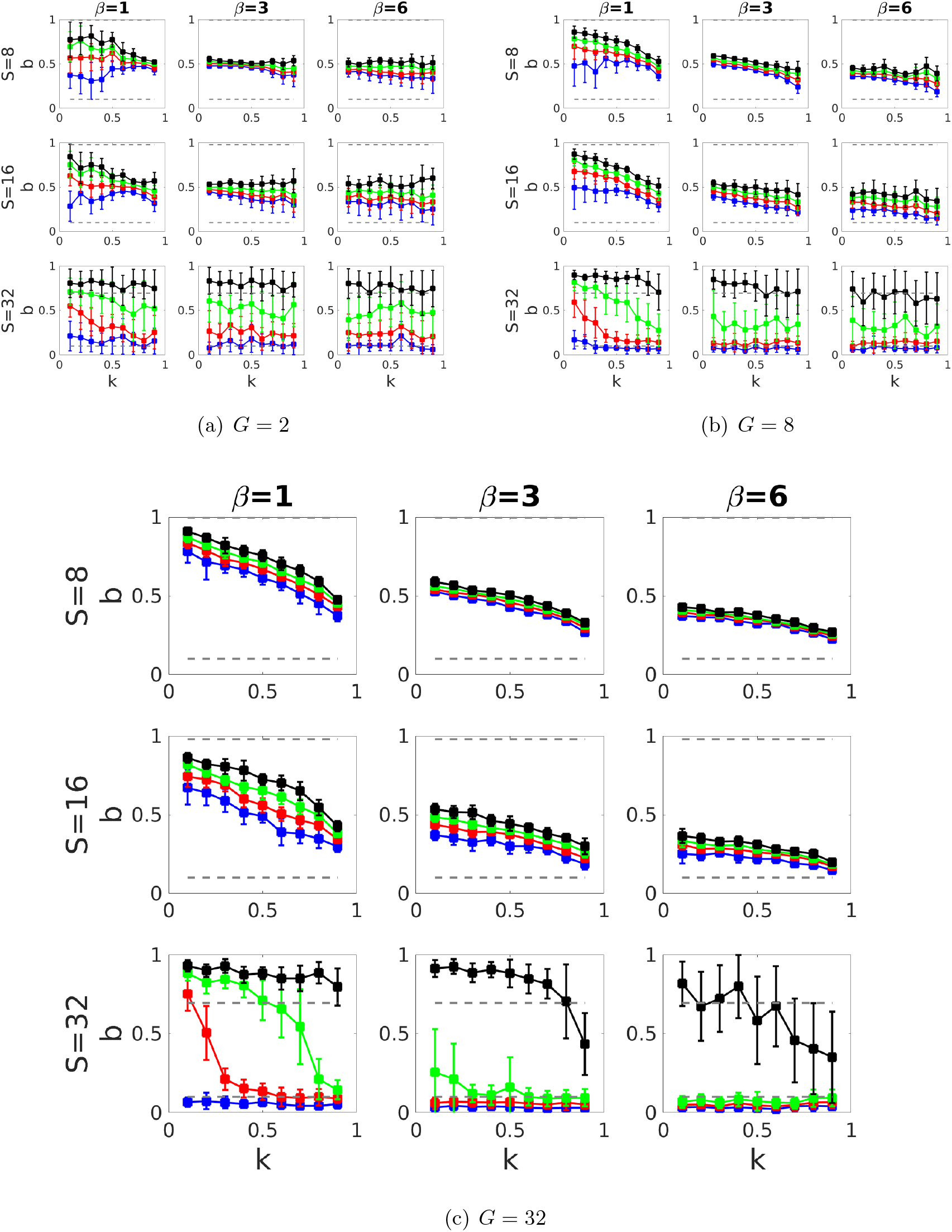
Effects of the relative cost of fecundity *k*, the strength of competition *β* and the number of cells *S* on fecundity *b* of different cell prototypes in the case of random microenvironmental effects. Prototypes are sorted by increasing fecundity and marked with different colors. Dashed lines show cell phenotypes with *b* = 0.1 and *a* = 0.1, respectively. Baseline parameters: *s* = 4, *C* = 1000, *ω* = 4, *µ* = 0.001 and *σ* = 0.05. Parameter *γ* = 2 for *S* = 8, *γ* = 3*/*2 for *S* = 16 and *γ* = 2*/*3 for *S* = 32. Shown are averages and confidence intervals based on 20 runs each of length 10^6^ time steps for each parameter combination. Results on each run have been calculated based on the last quarter of time steps.

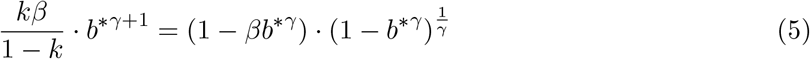

The above equation cannot be solved analytically in the general case. However, we can get some analytical progress for a number of special cases of the model (see details in the Supplementary modeling section in the SM). Moreover, one can show that increasing *k* or *β* decreases *b*^*\**^ for chosen model parameters.

If the number of genes is small (i.e *G* = 2 or 8), then more than one cell type per colony can be observed (Figure 2a,b and Figure S1a in the SM). Specifically, two or more cell phenotypes are observed if both *β* and *k* are simultaneously small or large. With rare exceptions, all these cell phenotypes are unspecialized (Figure S1b in the SM).

#### Large colonies

Following Michod (2005, 2006); Michod *et al.* (2006), we assume that in large colones (here with *S* = 32 cells), the trade-off function describing the relationship between activity and fecundity is convex with *γ <* 1.

Our analytical results show that for the large number of genes (*G* → ∞) two different cell phenotypes are present in the colony: there is a differentiation into somatic and reproductive cells, or the colony is composed by somatic and unspecialized cells (only if *β* and *k* are large). For small *β* the share of reproductive cells *p*_*g*_ decreases with an increase in *k*. For intermediate or large *β*, the majority of cells are somatic regardless of *k*. For more details see the Supplementary modeling section the SM.

The above results are supported by numerical simulations for the case of large number of genes (i.e., *G* = 32). Assume that *β* is small. Typically, a differentiation into somatic and reproductive cells within a colony is observed. For small *k* the majority of cells specialize in the reproductive function. Cell prototypes one-by-one switch to specialization in somatic function as *k* increases. The third, unspecialized cell type, can emerge during these transitions. For large *k* the majority of cells specialize in the somatic function. Assume that *β* is intermediate or large. Increasing *k* first has no effect on the equilibrium colony phenotype: it is composed by reproductive and somatic cells with higher number of somatic cells. For sufficiently large *k* reproductive cells disappear, and the colony is composed of somatic and unspecialized cells. For more details see Figure 2c and Figure S1 in the SM).

If the number of genes is small (i.e. *G* = 2 or 8), then on average a large number of distinct cell phenotypes is observed (see Figure 2a,b and Figure S1a in the SM). Moreover, decreasing the number of genes (e.g., from *G* = 8 to *G* = 2) increases the number of different cell phenotypes simultaneously reducing the share *p* of specialized cells (Figure S1a,b in the SM). For example, on average three different cell phenotypes are present, if there are just two genes (Figure S1a in the SM).

The strong effect of the number of genes *G* on the model dynamics can be explained in the following way. Let us define a fitness of a focal colony as an expected number of offspring colonies that are started from the focal one. Let us define a globally optimal colony phenotype as one that maximizes fitness subject to trade-off constraints and the resource constraint [recall that we define a colony phenotype as a set of all cell phenotypes (cell fecundities) in the colony]. Cell microenvironment imposes additional constraints, therefore some colony phenotypes become unattainable. According to Equation 1 an increase in the number of genes increases the number of ways a colony can interact with its environment, which in turn reduces the strength of microenvironmental constraints. As a result, increasing *G* increases the colony fitness. Moreover, it moves the equilibrium colony phenotype closer to the globally optimal one. With *G* → ∞ the globally optimal colony phenotype is attained.

### 3.2 Microenvironmental gradients

Similar to the previous section, in the case of microenvironmental gradients the system evolves to an equilibrium at which between-colony variation in colony phenotypes is very low. However, the cell phenotypes observed can be quite different from those in the case of random microenviron-mental effects. Note that models in this section have a new parameter: the decay rate in chemical concentration *r* along the anterior-posterior axis. We will consider the case of a single gradient (*E* = 1) separately.

#### 3.2.1 Single gradient

Here we will first discuss results that are observed for both small-size and large-size colonies, then we will provide results on small-size colonies, and finally results on large-size colonies.

In the case of a single gradient there is a key difference with the results reported so far. Namely, we can observe that the number of genes *G* has an insignificant impact on the model dynamics, regardless of the colony size *S* (see Figure 3 and Figures S2-S5 in the SM). However, the effect of parameters *β* and *k* on the population size *N*, the group competitiveness *A*, and the total fecundity *B* are similar to those observed with random microenvironmental effects. With rare exceptions, the decay rate *r* in chemical concentration has no effect on the above characteristics. The results are shown in Figure S5 in the SM.

**Figure 3:**
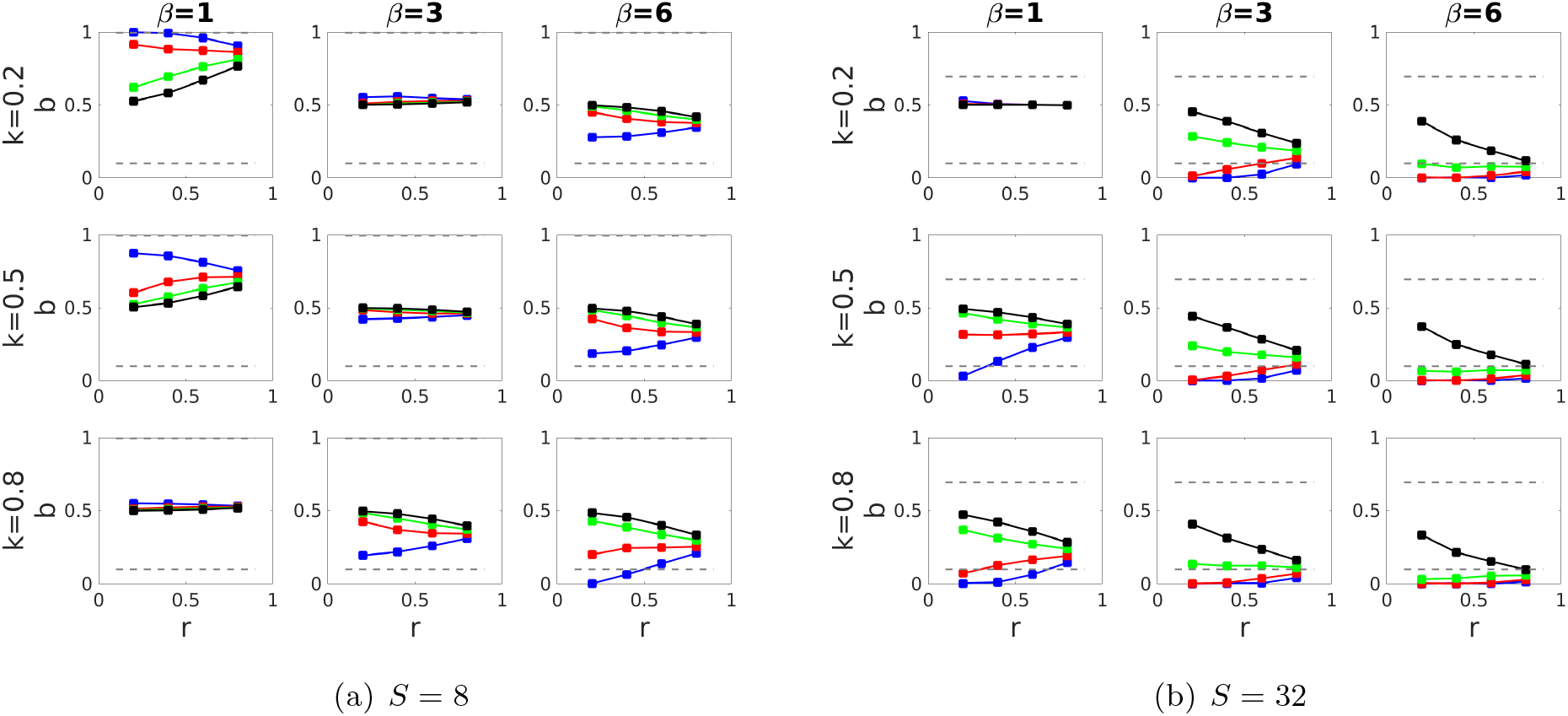
Effects of the relative cost of fecundity *k*, the strength of competition *β* and the decay rate *r* in chemical concentration on the equilibrium cell phenotypes *b* in the case of a single gradient. Cell prototypes are marked in blue, red, green and black colors respectively along the anterior-posterior axis. Dashed lines show cell phenotypes with *b* = 0.1 and *a* = 0.1, respectively. Baseline parameters: *G* = 32, *s* = 4, *d* = 10, *C* = 1000, *ω* = 4, *µ* = 0.001 and *σ* = 0.05. Shown are average results based on 20 runs each of length 10^6^ time steps for each parameter combination. Results on each run have been calculated based on the last quarter of time steps.

##### Small colonies

As it was discussed so far, a small-size colony obtains the maximum fitness if it is composed by identical unspecialized cells with fecundity *b*^*\**^ defined by Equation 5. This state can be unattainable due to microenvironmental constraints. The strength of these constraints is controlled by the decay rate *r* in chemical concentration. Specifically, differences in microenvironment experienced by distinct prototypes are subsequently amplified by decreasing *r*. As a result, for large *r* a colony is composed by identical unspecialized cells, while decreasing *r* leads to an increase in the number *M* of different cell phenotypes present in a colony (see Figure 3a and Figure S2a in the SM).

Some of these cell phenotypes are specialized. Namely, a colony comprises reproductive cells, if *k, β* and *r* are small. Somatic cells are present in a colony, if *k, β* are large and *r* is small. In all other cases only unspecialized cells are observed (these can involve different types of unspecialized cells). The results are shown in Figure S4a in the SM. Generally, specialized cells are located in the anterior layers (Figure 3a).

##### Large colonies

As it was discussed in the previous section, in general, a large-size colony obtains the maximum fitness, if there is a differentiation into somatic and reproductive cells. This outcome never arose with a single gradient (see Figure 3b and Figures S2b-S4b in the SM).

Different number *M* of cell phenotypes can be observed in the model. Specifically, one cell type is present in a colony, if (i) *r* is large or (ii) *k* and *β* are small. Otherwise from two to four distinct cell phenotypes are detected. In general, decreasing *r* increases *M* . The results are illustrated in Figure S2b in the SM.

Interestingly, reproductive cells never emerge while somatic cells are observed for a broad set of parameter values (see Figure 3b and Figure S4b in the SM). To explain why reproductive cells never emerge, note that in the case of a single gradient either activity is suppressed in all cells of a colony (i.e., cumulative gene effects *x* are positive for all cells), or fertility is suppressed in all cells of a colony (i.e., cumulative gene effects *x* are negative for all cells). According to Equation 1 along with the fact in the previous sentence it means that each cell in a colony has its fecundity either higher than 0.5 or lower than 0.5. The former case cannot be observed at equilibrium. Indeed, if each cell has fecundity higher than 0.5, too many offspring will be produced (since large-sized colonies are considered). This will intensify the resource-based competition. As a result, colonies that contribute more to activity will have higher fitness. Therefore, each cell in a colony has either (i) fecundity that is lower than 0.5 (general case), or (ii) fecundity that is a little bit higher than 0.5 (the case of small *k* and *β*).

As it is shown in Figure S4b, increasing the strength of competition *β* as well as the relative cost of fecundity *k*, promote the emergence of somatic cells. Up to 75% of cells can be identified as somatic if *β* and *k* are large.

Overall, we conclude that an equilibrium colony phenotype observed with a single gradient can be close to the globally optimal phenotype in the case of small colonies, and it is displaced from the globally optimal phenotype in the case of large colonies.

#### 3.2.2 Several gradients

##### Small colonies

All the results are similar to those observed with random microenvironmental effects and large number of genes. Specifically, at equilibrium, all cells in the colony have the same unspecialized phenotype. This phenotype is very close to the globally optimal one, regardless of the number of genes *G*. For more details see Figure 4 and Figures S6-S7 in the SM.

**Figure 4:**
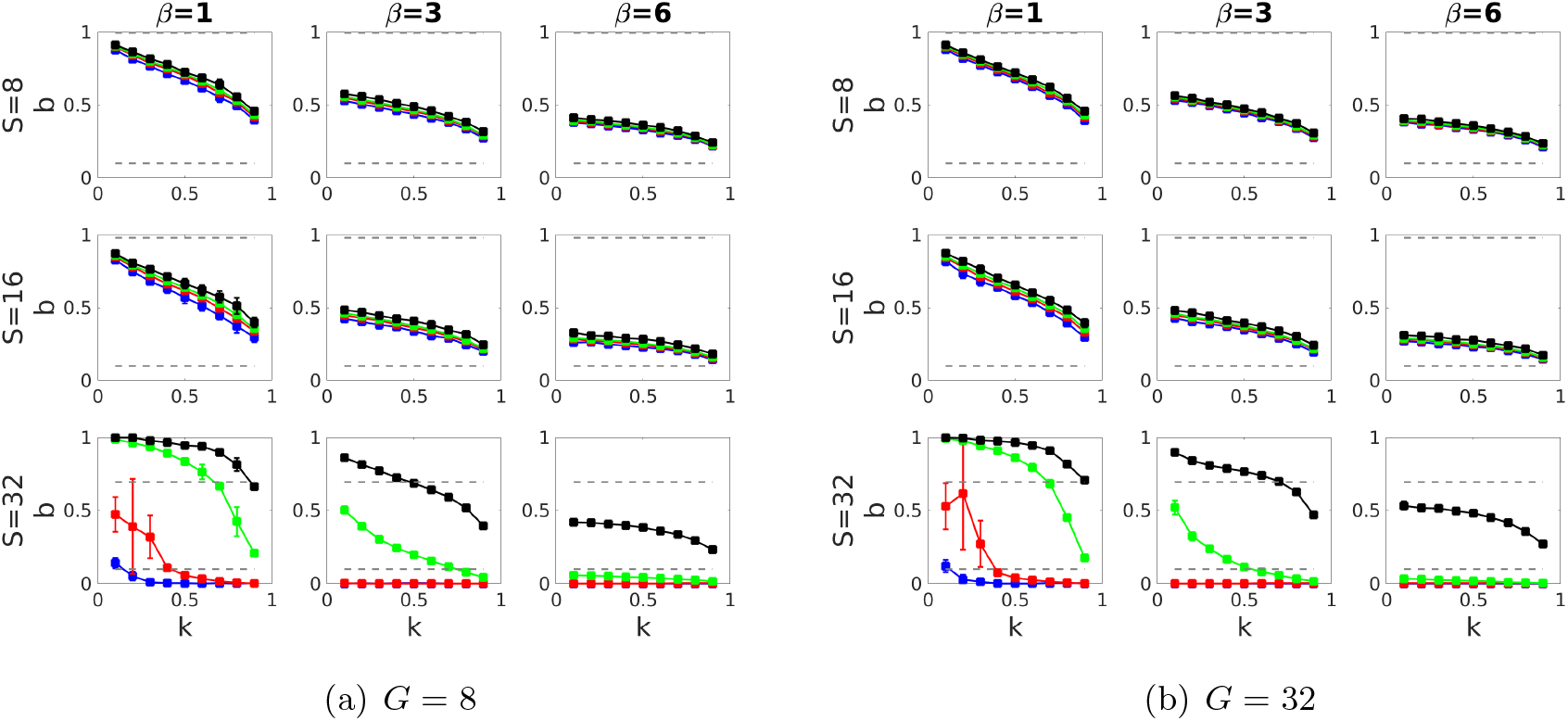
Effects of the relative cost of fecundity *k*, the strength of competition *β* and the number of cells *S* on fecundity *b* of different cell prototypes in the case of *E* = 4 different microenvironmental gradients. Prototypes are sorted by increasing fecundity and marked with different colors. Dashed lines show cell phenotypes with *b* = 0.1 and *a* = 0.1, respectively. Baseline parameters: *s* = 4, *C* = 1000, *ω* = 4, *µ* = 0.001 and *σ* = 0.05. Parameter *γ* = 2 for *S* = 8, *γ* = 3*/*2 for *S* = 16 and *γ* = 2*/*3 for *S* = 32. We assume that *r*1 = 0.2, *r*2 = 0.4, *r*3 = 0.6 and *r*4 = 0.8 for the first, second, third and fourth gradients respectively. Shown are averages and confidence intervals based on 20 runs each of length 10^6^ time steps for each parameter combination. Results on each run have been calculated based on the last quarter of time steps.

##### Large colonies

All the results are close to those observed with random microenvironmental effects and large number of genes (see Figure 4 and Figures S6-S7 in the SM). Typically, a colony is composed by two cell phenotypes: (i) reproductive and somatic, or (ii) unspecialized and somatic; or three cell phenotypes are present: (iii) somatic, reproductive and unspecialized (Figure 4 and Figure S6 in the SM). The former outcome is less likely observed compared to the case of random microenvironmental effects.

If *β* and *k* are small, the colony is composed by the three cell phenotypes: somatic, reproductive, and unspecialized (with half of the cells specialized in reproductive function). If *β* is small and *k* is intermediate, cells in a colony are differentiated into somatic and reproductive. Increasing *β* or *k* decreases the total fecundity of a colony. As a result, at least half of the cells in a colony are identified as somatic for intermediate or large values of *β* or large values of *k*. Eventually, for large *β* a colony is composed by somatic and unspecialized cells, and somatic cells predominate. The number of genes *G* has a limited effect on the model dynamics. Increasing *G* reduces the number of different cell phenotypes *M* increasing the share of specialized cells *p* in a colony, if *β* is intermediate; and has no significant effects otherwise. For more details see Figure 4 and Figure S6 in the SM

Overall one can conclude that the results obtained for several gradients differ significantly from those observed for a single gradient. Thus, an equilibrium colony phenotype becomes much closer to the globally optimal one compared to the case of a single gradient. Overall, we conclude that an environment with several gradients brings more fitness advantages to a colony compared to one with just a single gradient.

## 4 Discussion

Division of labor can be found at different levels of biological organization - from genes and cell colonies to eusocial insects and human societies (Smith 2008; Rodriguez-Clare 1996; Stigler 1951; Robinson 1992; Fay 1992; Panter-Brick 2002; Berman-Frank *et al.* 2003; Kirk 2003; Nedelcu and Michod 2006; Flores and Herrero 2010; Rossetti *et al.* 2010; Reuffler and Wagner 2012; Duarte *et al.* 2011; Durkheim 2014). One of the simplest examples of the division of labor in multicellular organisms is germ-soma specialization, which plays a key role in the evolution of organismal complexity (Smith and Szathmary 1995). Here we have formulated and studied a general mathematical model exploring the emergence of germ-soma specialization in colonies of cells. We have considered a finite population of colonies competing for resources. In our model, colonies are of the same size and are composed by asexually reproducing haploid cells. Each cell can contribute to activity and fecundity of the colony, these contributions are traded-off. An amount of the resource a colony secured depends on its relative activity, and determines its probability of survival. We have assumed that all cells within a colony are genetically identical but gene expression is affected by variation in microenvironment experienced by individual cells. We have found conditions that favour the emergence of germ-soma specialization under different types of microenvironmental effects. Specifically, we have explored the case of random microenvironmental effects and the case of microenvironmental gradients. In the latter case, gene expression levels change in accordance with some chemical gradients acting on the colony. The results of our modeling are summarized in Table 1. Below we discuss our major findings as well as their biological interpretation.

**Table 1:** The main results.

| Micro-environment | Colony size | Number of genes | Typical patterns of within-colony differentiation | Conditions for coexistence of reproductive and somatic cells |
| --- | --- | --- | --- | --- |
| Random | Small | Large | One type of unspecialized cells | - |
|  |  | Small | One or two types of unspecialized cells | - |
| | Large | Large | Two types of cells: somatic and reproductive or somatic and unspecialized | All cases except one with large $k$ and $\beta$ |
| | | Small | Three types of cells: somatic, reproductive and unspecialized | Can be observed for all $k$ and $\beta$, conditions are determined by random microenvironmental effects |
| Single gradient | Small | Not important | One type of unspecialized cells, or 2 types of cells (somatic and unspecialized or reproductive and unspecialized) | - |
|  | Large | - " - | One type of unspecialized cells, or 2 types of cells (somatic and unspecialized or 2 unspecialized) or 3 types of cells (somatic and 2 unspecialized) | - |
| Several gradients | Small | - " - | One type of unspecialized cells | - |
| | Large | - " - | Two types of cells (somatic and reproductive or somatic and unspecialized) or three types of cells (somatic, reproductive and unspecialized) | $\beta$ is not large and $k$ is not large |

We have shown that the shape of the trade-off relationship between cells’ fecundity and activity is a crucial factor controlling the overall proportion of specialized cells (somatic or reproductive) within the colony. We have found that larger convexity of the trade-off relationship (expected in larger colonies) increases the share of specialized cells. This is well in line with the current literature (Egas *et al.* 2004; Michod 2005, 2006; Michod *et al.* 2006; Reuffler and Wagner 2012; West *et al.* 2015; Cooper and West 2018). However we have found that specialization can evolve even if the curvature of the trade-off relationship is concave. Moreover our model predicts that incomplete specialization or even colonies composed only by unspecialized cells can be observed in the case of a convex trade-off. This phenomenon has already been reported in the literature. For example, Yanni *et al.* (2020) considered topological structures of interactions between colony cells and demonstrated that germ-soma specialization can emerge for a broad class of network topologies even when the trade-off relationship is concave. Tverskoi *et al.* (2018) explored the effects of resource constraints on the differentiation of somatic and reproductive cells. They showed that a strong resource constraint can promote the emergence of division of labor in colonies with concave trade-offs as well as lead to the collapse of specialization in colonies with convex trade-offs. In our model, this phenomenon arises as a consequence of certain microenvironmental effects experienced by cells. Specifically, specialization can evolve when the trade-off relationship is concave in the case of a single environmental gradient. Unspecialized cells can be observed in colonies with convex trade-offs in the case of a single gradient or in the case of random microenvironmental effects and a small number of genes.

We have demonstrated that the type and extent of variation in within-colony microenvironment and, in the case of random microenvironment, the number of genes involved, are key factors shaping the model dynamics and, hence, the emergence of germ-soma separation. In the case of random microenvironmental effects, if the number of genes and the colony size are large, there is differentiation into somatic and reproductive cells or the colony is composed by somatic and unspecialized cells. Decreasing the number of genes reduces the share of specialized cells in large colonies. We have shown that in general, if the colony size is small, only unspecialized cells are present. Overall, increasing the number of genes reduces genetic constraints faced by cells and increases the colony fitness (i.e., the expected number of offspring colonies). Similar effects can be found in some experimental works. For example, a large share of all genes detected in the yeast genome have neither established nor acquired function (Goebl and Petes 1986; Oliver *et al.* 1992). According to the marginal benefit hypothesis these genes are not essential, but instead they make small contributions to the efficiency of various processes occurring within a colony. As shown in Thatcher *et al.* (1998), mutants with some non-essential genes suppressed (in terms of our model these mutants can be treated as ones with a lower number of genes) have a significantly lower fitness compared to non-mutant individuals.

In the case of microenvironmental gradients, if their number and the colony size are large, there is differentiation into somatic and reproductive cells or the colony is composed by somatic and unspecialized cells. The former outcome can also involve some unspecialized cells. If the number of gradients is large and the colony size is small, only unspecialized cells are present. If there is only one microenvironmental gradient, all cells are unspecialized or the colony is composed by somatic and unspecialized cells (small and large colonies) or the colony is composed by reproductive and unspecialized cells (only small colonies). Increasing the number of gradients also increases the colony fitness. This happens because increasing the number of environmental gradients leads to an increase in the dimensionality of phenotype space. This in turn generates a fitness landscape with a new fitness maximum, in which colony phenotypes that are fitness optimal for smaller number of gradients become fitness saddle points. This effect of increasing number of gradients is analogous to the effects of cell aggregation on the emergence of cell specialization (Ispolatov *et al.* 2012).

We have shown that in our model the share of reproductive cells in large groups is relatively small, with the exceptions of some special cases characterized by a small strength of competition *β* and a small relative cost of fecundity *k*. This result is intuitive: a mutant colony with a large number of reproductive cells produces a large number of offspring. Due to the trade-off constraints, these offspring colonies have low activity, resulting in a very low probability of survival in the resource-based competition. This effect is amplified by increasing the strength of competition *β*, the relative cost of fecundity *k* and the colony size *S*. Eventually, it will lead to the extinction of the mutant phenotype. This outcome can be illustrated by volvocine green algae biology, where the largest forms (like various Volvox species) consist mainly of somatic cells and a small number of reproductive cells (Herron and Michod 2007; Shelton *et al.* 2012).

Several additional points and connections are worth noting. First, as it was shown in Amado *et al.* (2017, 2018); Yanni *et al.* (2020), the geometry of multicellular organisms can significantly influence the evolution of cell specialization. In our model, the group geometry is implicitly captured by different microenvironmental conditions experienced by cells of different prototypes (where a prototype is characterized by cells sharing the same vector of microenvironmental effects). However the number of different prototypes *s* reflecting the geometry of a colony was determined endogenously. Potentially, our approach can be expanded to study a co-evolution of the colony geometry (i.e., the number of different cell prototypes *s* and the difference in chemical concentration along gradients experienced by cells of different prototypes) and germ-soma specialization. Second, in our model only two tasks, reproductive and somatic, were taken into account. A reasonable extension would be to consider multiple tasks (e.g., reproductive and several somatic). This will allow us to gain new insights into the emergence of multiple specialized cell phenotypes and theoretical foundations of the size-complexity rule (Bonner 2004). Third, our results are based on very simple assumptions about the colony life-cycle and its reproductive mode. It would be a worthwhile to investigate how the results change for colonies with more complex life-cycles (Pichugin *et al.* 2017; Staps *et al.* 2019; Pichugin *et al.* 2019; Pichugin and Traulsen 2020). We leave these extensions and generalizations for future work.

Overall, our paper provides an additional argument for the importance of environmental factors in the evolution of germ-soma separation and the emergence of a new level of biological organization - a multicellular organism with complete germ-soma differentiation.

## Acknowledgment

We thank S. Carrignon, A. Senthilnathan, and L. Santana Souza for comments and suggestions. The paper was prepared within the framework of the HSE University Basic Research Program and funded by the Russian Academic Excellence Project ‘5-100’. The paper was partially supported by the Office of Naval Research grant W911NF-17-1-0150. D.T. also thanks the International Center of Decision Choice and Analysis (DeCAn Center) of the National Research University Higher School of Economics.

## 5 Supplementary materials

- Part 1 provides more results of numerical simulations underlying our conclusions presented in the main text. Figures are structured according to their links to the subsections of the main text.
- Part 2 provides detailed analytical results on globally optimal colony phenotypes.

### 5.1 Additional simulation results

#### 5.1.1 Random microenvironmental effects

**Figure S1:**
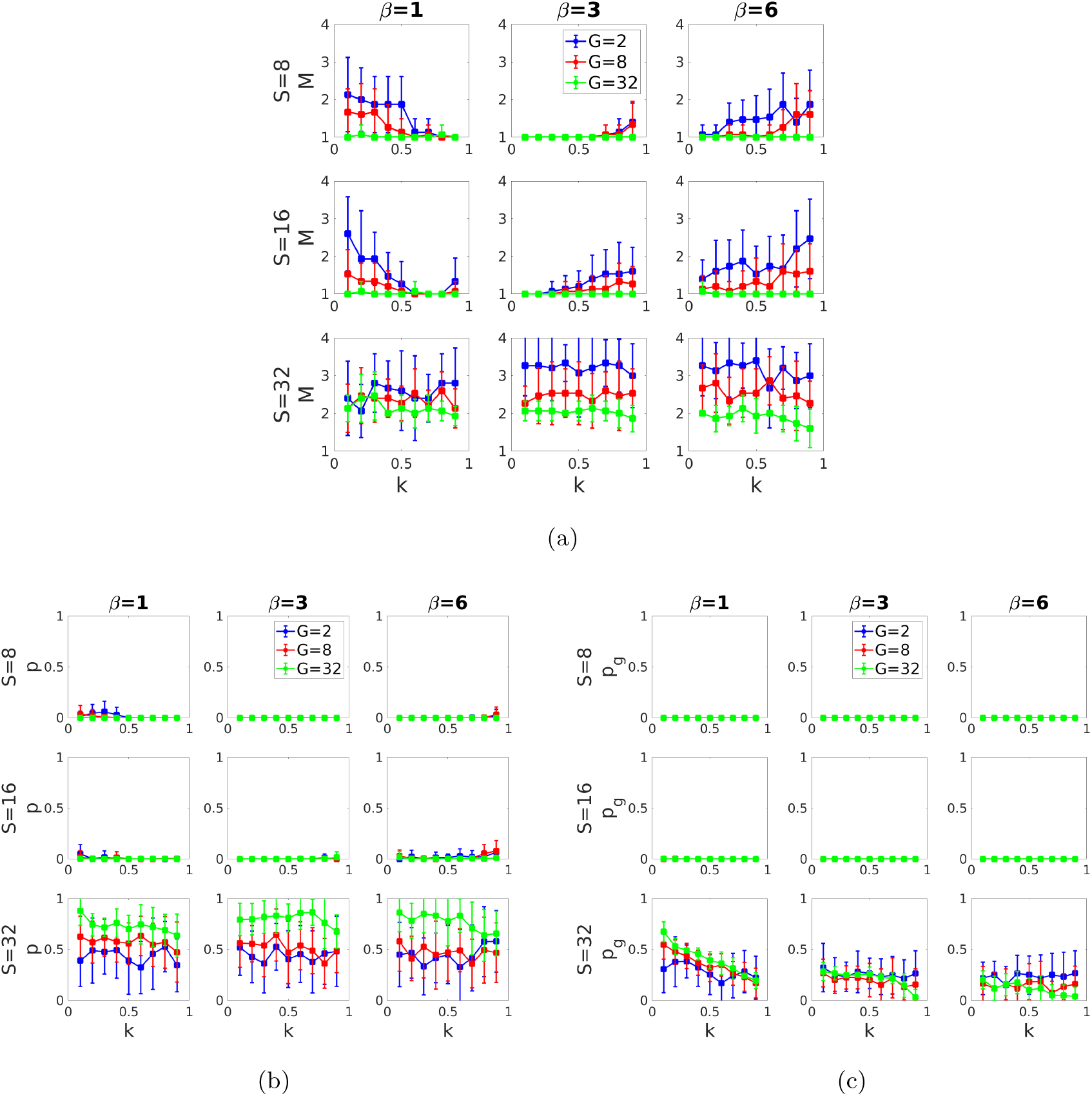
Effects of the relative cost of fecundity *k*, the strength of competition *β*, the number of cells *S* and the number of genes *G* on (a) the number of different cell phenotypes *M*, (b) the share *p* of specialized cells, and (c) the share of reproductive cells in a colony in the case of random microenvironmental effects. Baseline parameters: *s* = 4, *C* = 1000, *ω* = 4, *µ* = 0.001 and *σ* = 0.05. Parameter *γ* = 2 for *S* = 8, *γ* = 3*/*2 for *S* = 16 and *γ* = 2*/*3 for *S* = 32. Shown are averages and confidence intervals based on 20 runs each of length 10^6^ time steps for each parameter combination. Results on each run have been calculated based on the last quarter of time steps.

#### 5.1.2 Single gradient

**Figure S2:**
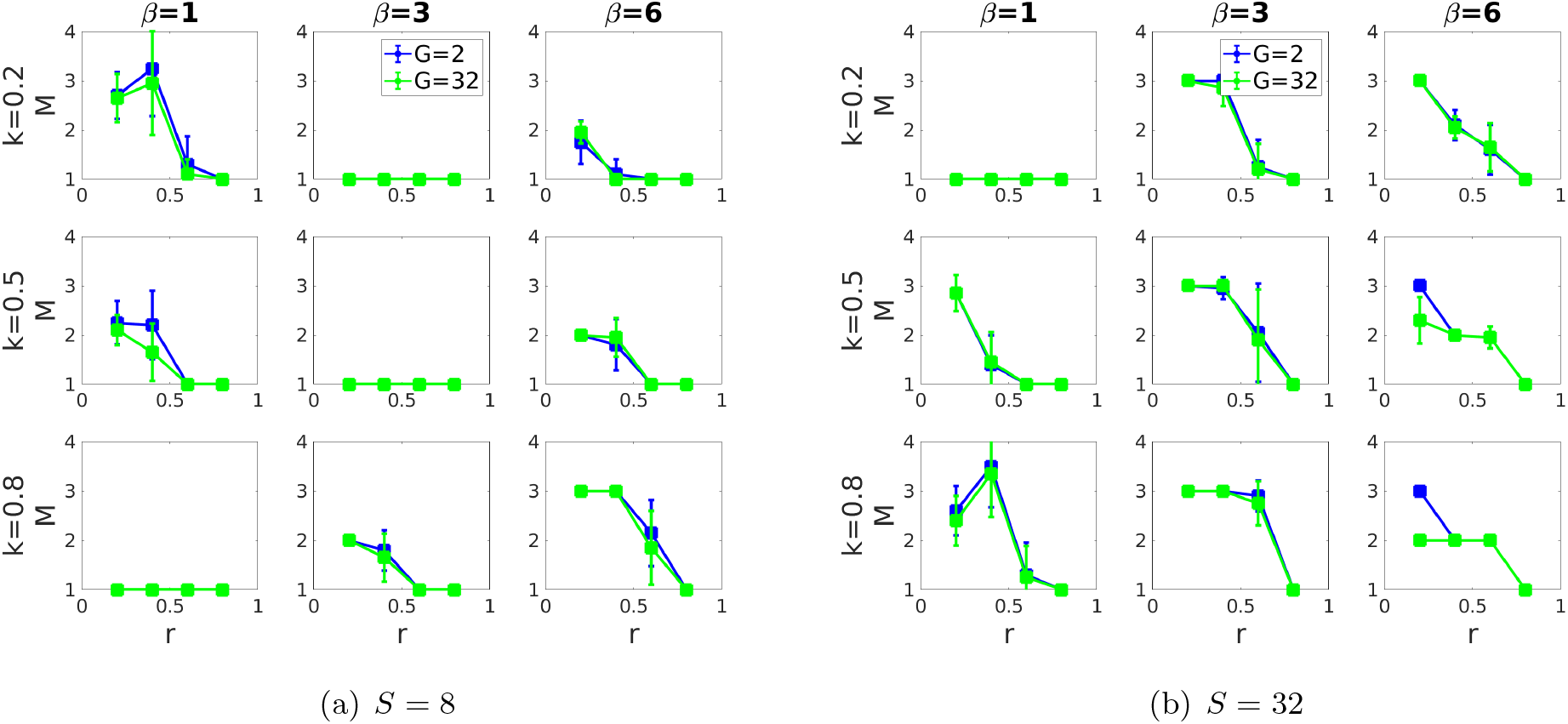
Effects of the relative cost of fecundity *k*, the strength of competition *β*, the decay rate *r* in chemical concentration, and the number of genes *G* on the number of different cell phenotypes *M* in a colony in the case of a single gradient. Baseline parameters: *s* = 4, *d* = 10, *C* = 1000, *ω* = 4, *µ* = 0.001 and *σ* = 0.05. Shown are average results based on 20 runs each of length 10^6^ time steps for each parameter combination. Results on each run have been calculated based on the last quarter of time steps.

**Figure S3:**
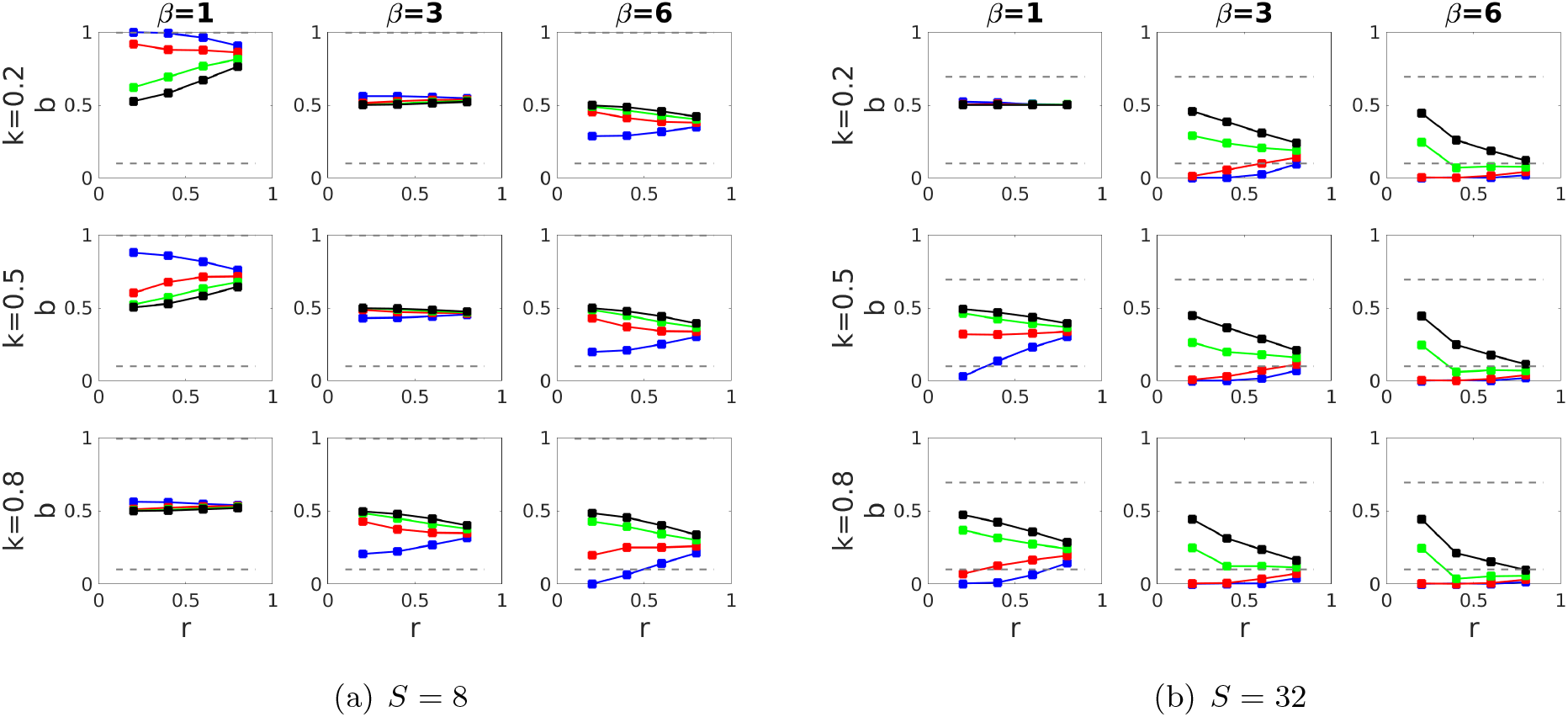
Effects of the relative cost of fecundity *k*, the strength of competition *β* and the decay rate *r* in chemical concentration on the equilibrium cell phenotypes *b* in the case of a single gradient. Cell prototypes are marked in blue, red, green and black colors respectively along the anterior-posterior axis. Dashed lines show cell phenotypes with *b* = 0.1 and *a* = 0.1, respectively. Baseline parameters: *G* = 2, *s* = 4, *d* = 10, *C* = 1000, *ω* = 4, *µ* = 0.001 and *σ* = 0.05. Shown are average results based on 20 runs each of length 10^6^ time steps for each parameter combination. Results on each run have been calculated based on the last quarter of time steps.

**Figure S4:**
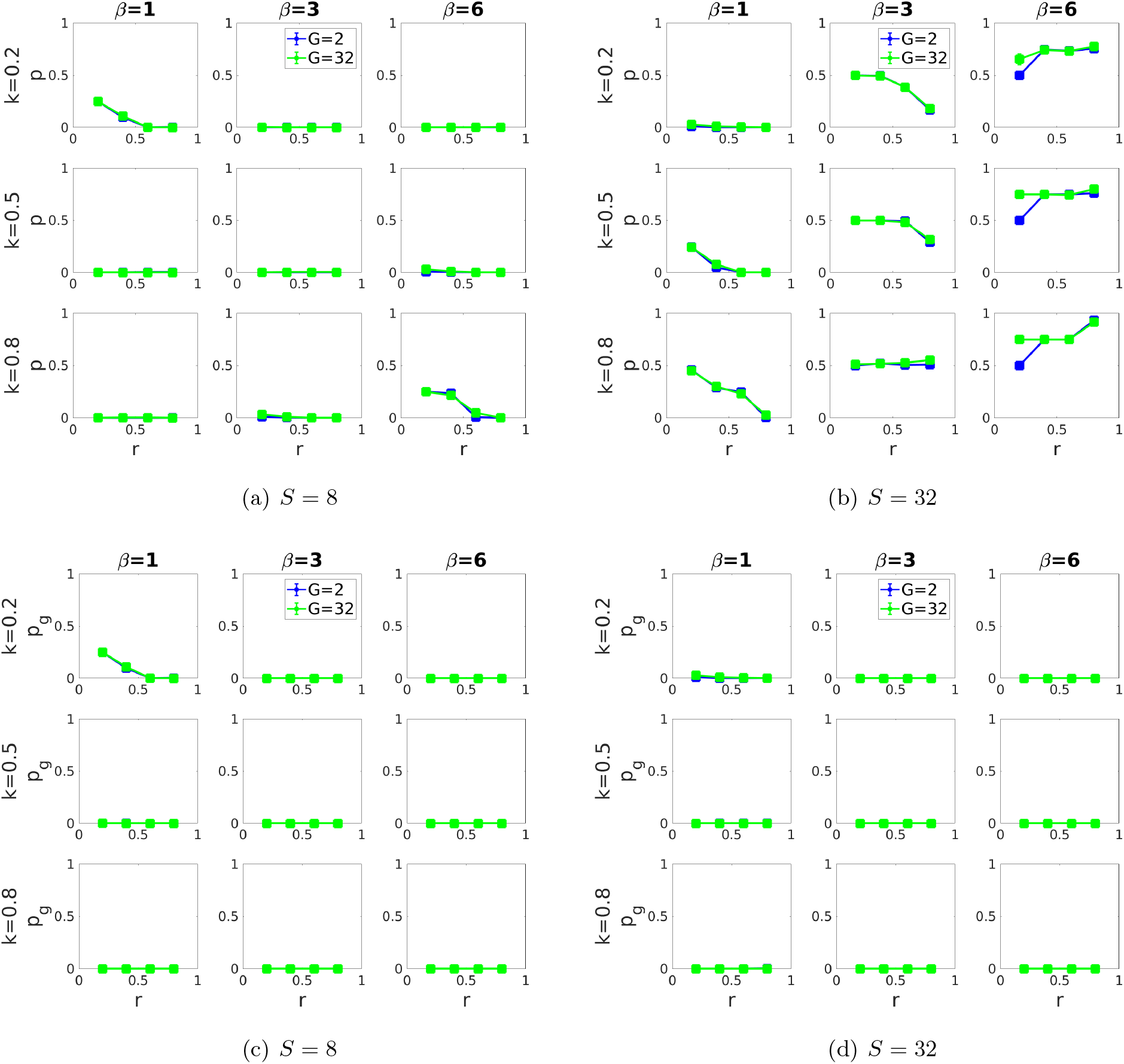
Effects of the relative cost of fecundity *k*, the strength of competition *β*, the decay rate *r* in chemical concentration, and the number of genes *G* on (a-b) the share *p* of specialized cells, and (c-d) on the share *pg* of reproductive cells in a colony in the case of a single gradient. Baseline parameters: *s* = 4, *d* = 10, *C* = 1000, *ω* = 4, *µ* = 0.001 and *σ* = 0.05. Shown are average results based on 20 runs each of length 10^6^ time steps for each parameter combination. Results on each run have been calculated based on the last quarter of time steps.

**Figure S5:**
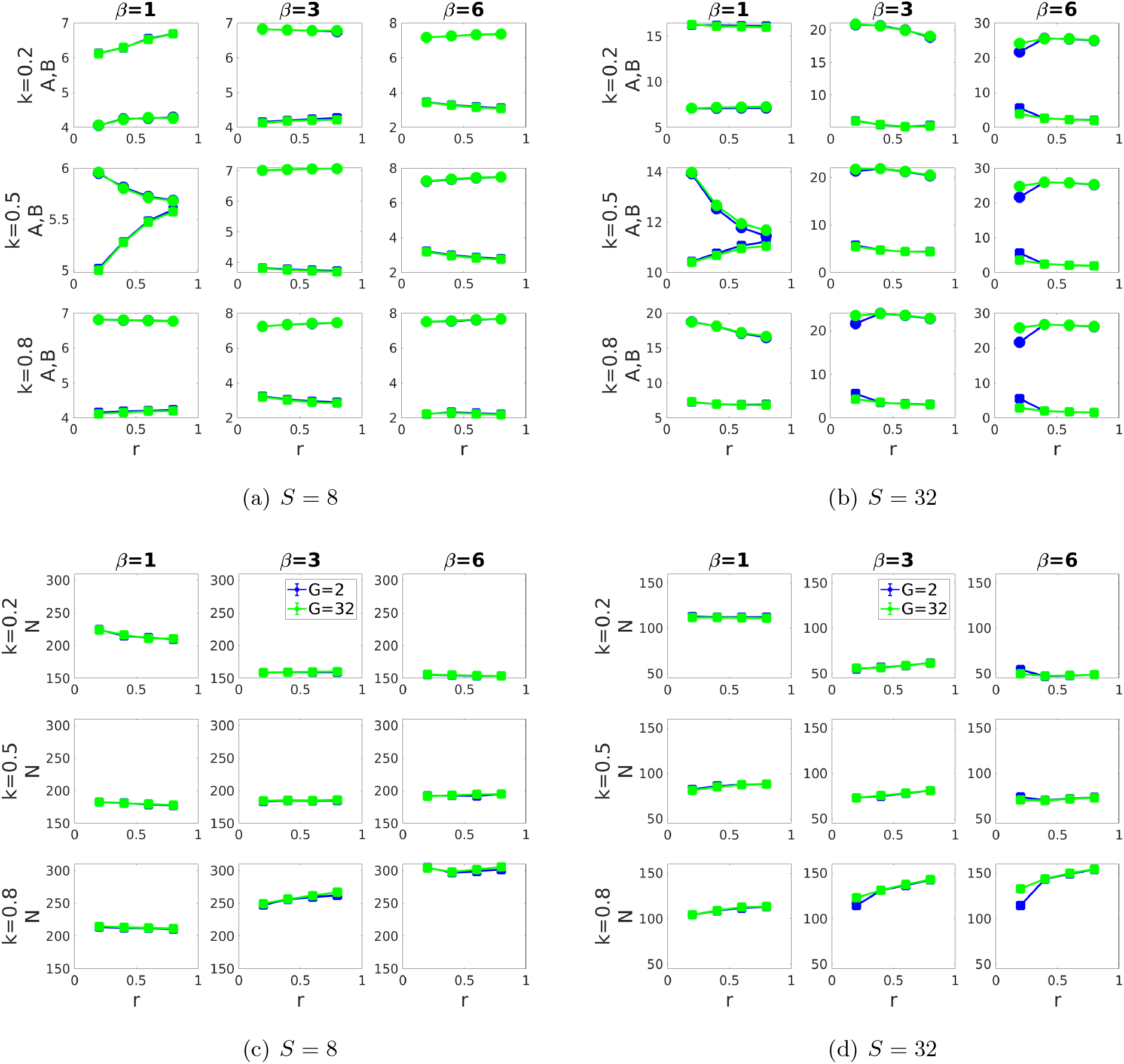
Effects of the relative cost of fecundity *k*, the strength of competition *β*, the decay rate *r* in chemical concentration, and the number of genes *G* on (a-b) the competitiveness *A* and the total fecundity *B* of a colony, and (c-d) the population size *N* in the case of a single gradient. Baseline parameters: *s* = 4, *d* = 10, *C* = 1000, *ω* = 4, *µ* = 0.001 and *σ* = 0.05. Shown are average results based on 20 runs each of length 10^6^ time steps for each parameter combination. Results on each run have been calculated based on the last quarter of time steps.

#### 5.1.3 Several gradients

**Figure S6:**
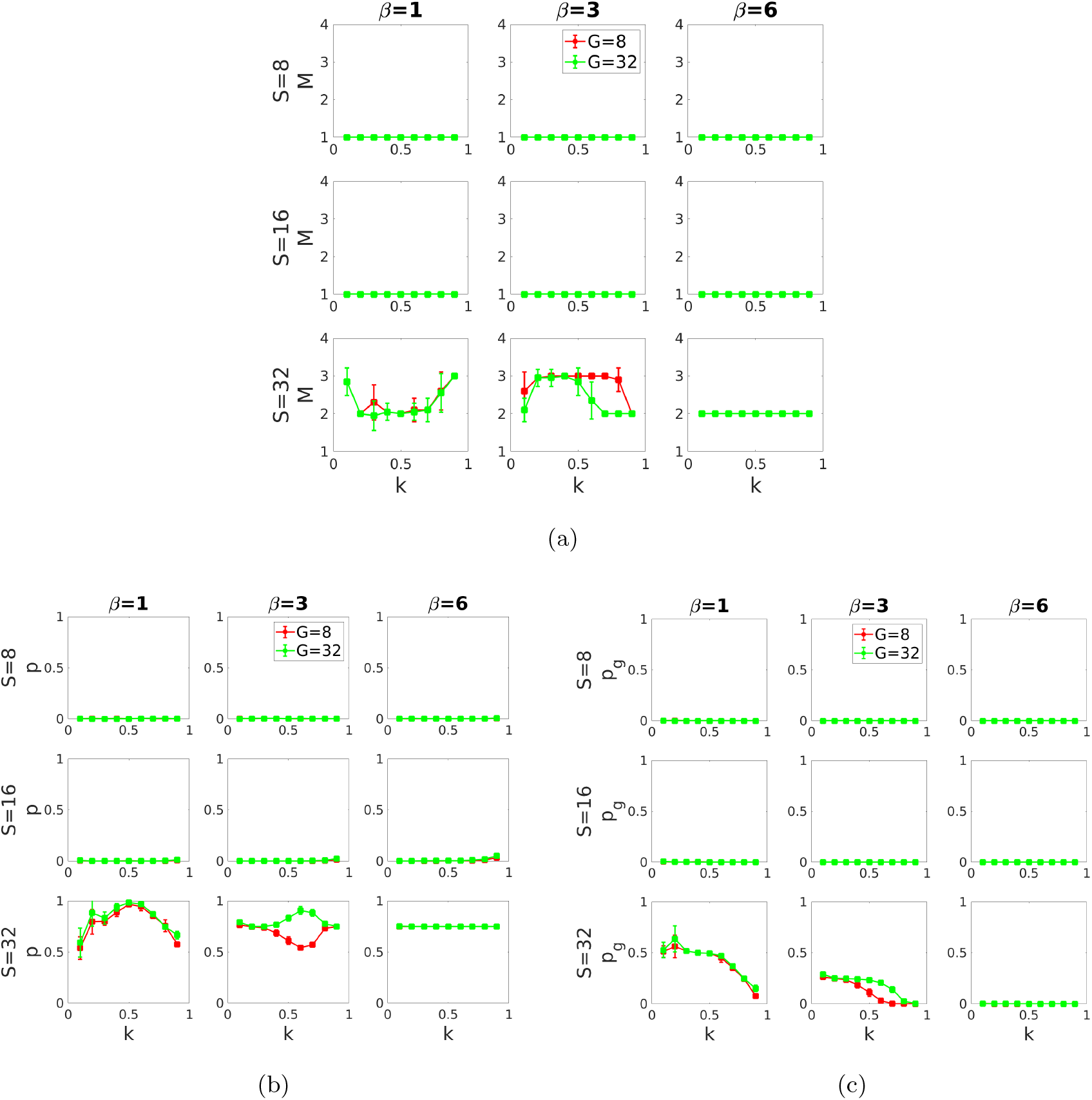
Effects of the relative cost of fecundity *k*, the strength of competition *β*, the number of cells *S* and the number of genes *G* on (a) the number of different cell phenotypes *M*, (b) the share *p* of specialized cells, and (c) the share of reproductive cells in a colony in the case of *E* = 4 different microenvironmental gradients. Baseline parameters: *s* = 4, *C* = 1000, *ω* = 4, *µ* = 0.001 and *σ* = 0.05. Parameter *γ* = 2 for *S* = 8, *γ* = 3*/*2 for *S* = 16 and *γ* = 2*/*3 for *S* = 32. We assume that *r*1 = 0.2, *r*2 = 0.4, *r*3 = 0.6 and *r*4 = 0.8 for the first, second, third and fourth gradients respectively. Shown are averages and confidence intervals based on 20 runs each of length 10^6^ time steps for each parameter combination. Results on each run have been calculated based on the last quarter of time steps.

**Figure S7:**
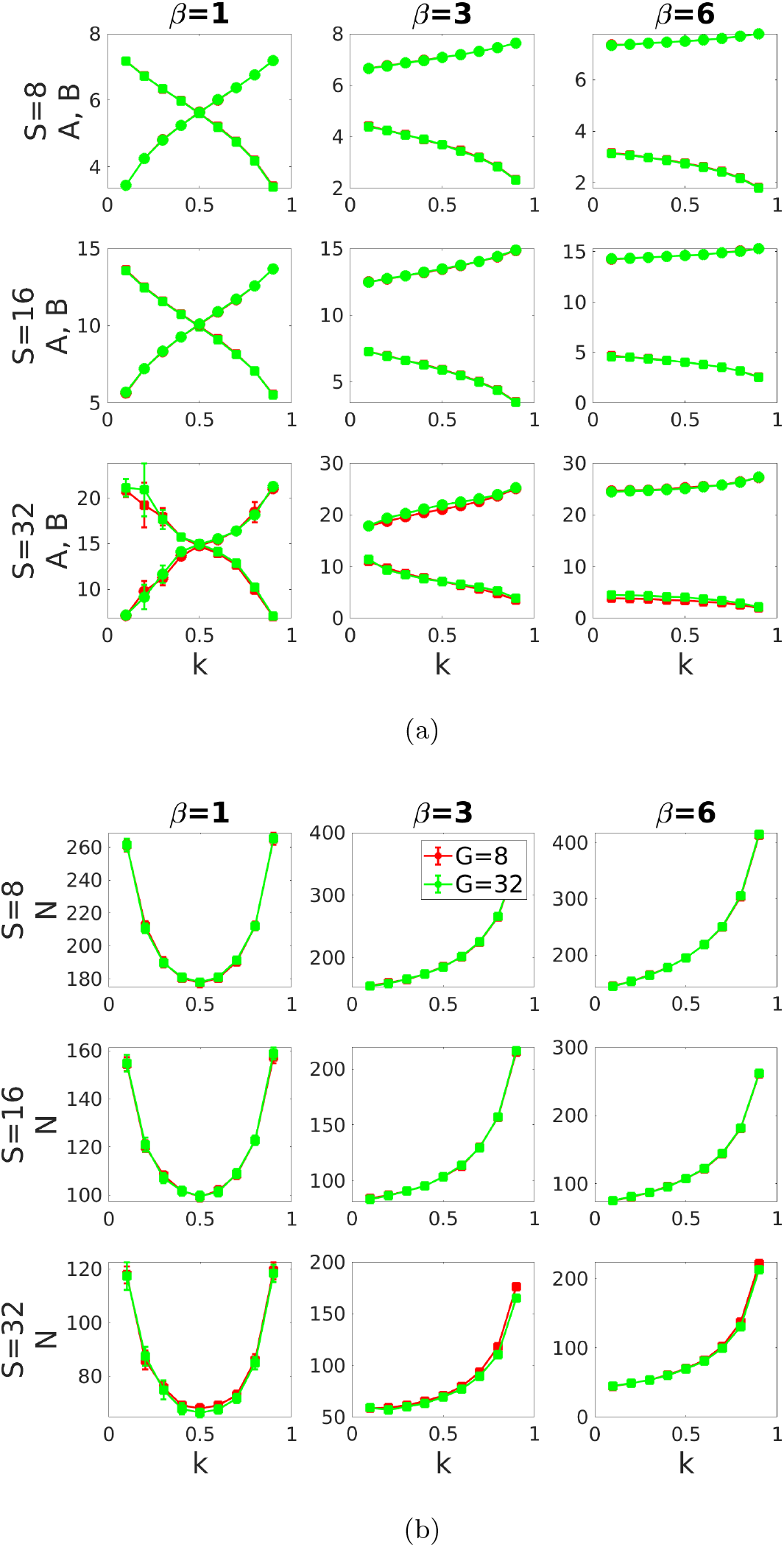
Effects of the relative cost of fecundity *k*, the strength of competition *β*, the number of cells *S* and the number of genes *G* on (a) the competitiveness *A* and the total fecundity *B* of a colony, and (b) the population size *N* in the case of *E* = 4 different microenvironmental gradients. Baseline parameters: *s* = 4, *C* = 1000, *ω* = 4, *µ* = 0.001 and *σ* = 0.05. Parameter *γ* = 2 for *S* = 8, *γ* = 3*/*2 for *S* = 16 and *γ* = 2*/*3 for *S* = 32. We assume that *r*1 = 0.2, *r*2 = 0.4, *r*3 = 0.6 and *r*4 = 0.8 for the first, second, third and fourth gradients respectively. Shown are averages and confidence intervals based on 20 runs each of length 10^6^ time steps for each parameter combination. Results on each run have been calculated based on the last quarter of time steps.

### 5.2 Supplementary modeling

Here we consider a finite population of colonies each composed by S asexually reproducing haploid cells with clonal development. Each colony is composed by cells of *s* different prototypes (1 ≤ *s* ≤ *S*). We postulate that there are *S*_*m*_ cells of prototype *m* in each colony, where *m* ∈ {1, .., *s*}. Cells of the same prototype have the same levels of fecundity and activity. We postulate that *β* ≥ 1 i.e., we only take into account those populations that live in a sufficiently competitive environment. For the sake of simplicity we will assume random microenvironmental effects.

#### 5.2.1 Fitness of a colony

A fitness of a new daughter colony in a resident population is defined as an expected number of offspring colonies that start from the daughter colony:

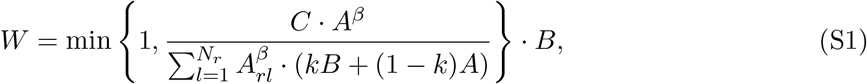

where the index *r* corresponds to colonies in the resident population. Since a new daughter colony will only reach reproductive age if it survives during the resource-based competition, there are two terms in the above formula. The first one, min 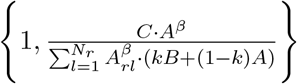, is equal to the probability of survival in the resource-based competition. The second one, *B*, is the expected number of offspring colonies that start from the adult one.

Let us define a globally optimal colony phenotype as one that maximizes the above fitness-function subject to trade-off constraints [recall that we define a colony phenotype as a set of all cell phenotypes (cell fecundities) in the colony]. Cell microenvironment imposes additional constraints on the phenotype space, therefore some colony phenotypes become unattainable. According to Equation 1 an increase in the number of genes increases the number of ways a colony can interact with its environment, which in turn reduces the strength of microenvironmental constraints. As a result, increasing *G* increases the colony fitness. Moreover, it moves the equilibrium colony phenotype closer to the globally optimal one. With *G* → ∞ the globally optimal colony phenotype is attained.

Let *B*^*\**^ and *A*^*\**^ be optimal values of the total fecundity and the total competitiveness of a colony, respectively. Then, the optimal value of the fitness function is

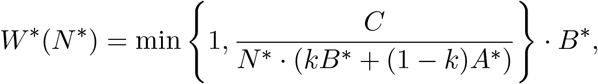

which reach a maximum, if the number of adult colonies *N* ^*\**^ is

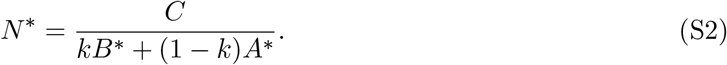

Below we present analytical results on the globally optimal colony phenotype for the case of small colonies and for the case of large colonies separately.

**Figure S8:**
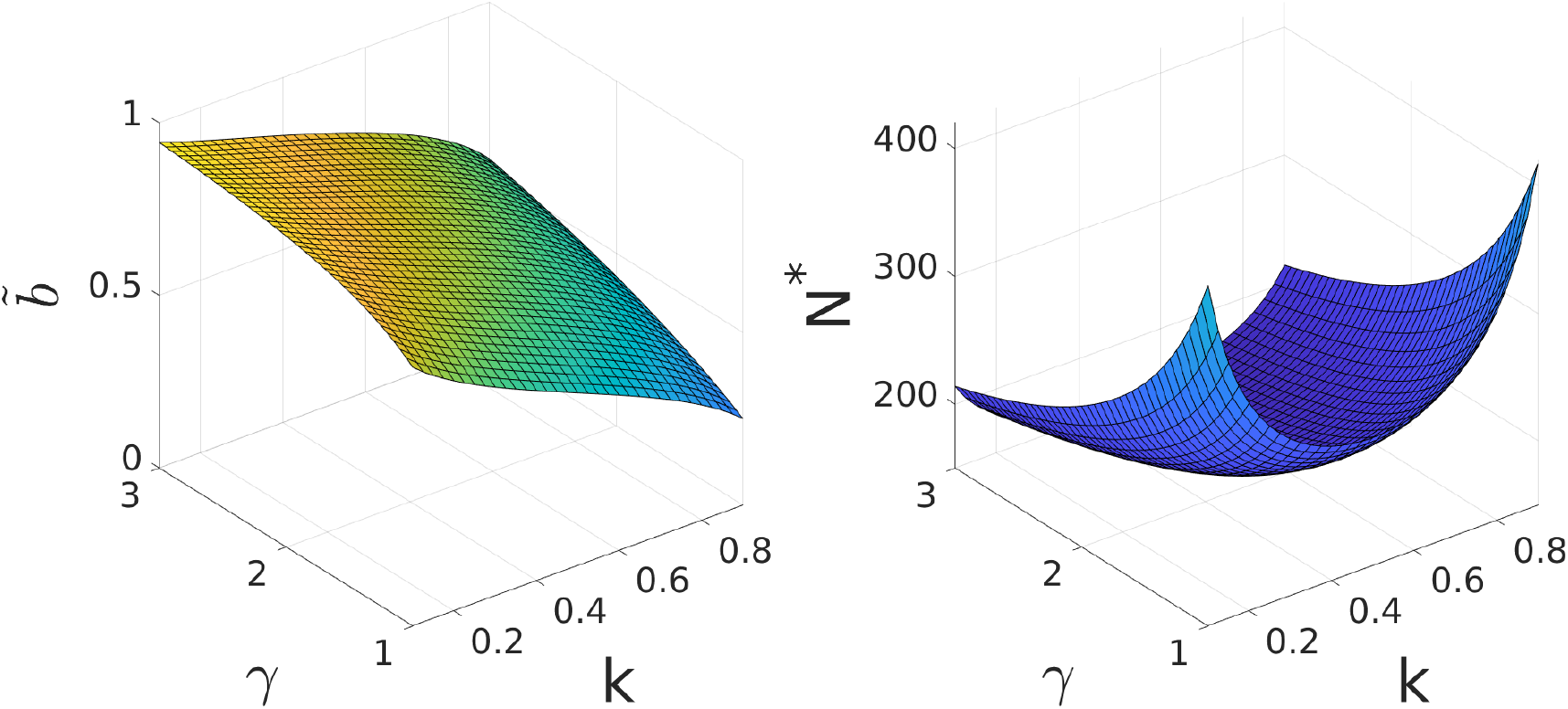
Analytical results for small colonies, *G* → ∞ and *β* = 1. Effects of the relative costs of fecundity *k* and the shape of the trade-off relation *γ* on the equilibrium cell phenotype 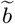 (the left panel) and the equilibrium population size *N* ^*\**^ (the right panel). Baseline parameter: *C* = 1000, *S* = 8.

#### 5.2.2 Fitness landscape analysis for a population of small colonies

Here we assume that in small colonies the trade-off function describing the relationship between activity *a* and fecundity *b* is concave with *γ* > 1.

Since cells of the same prototype have the same fecundity, the domain of the fitness function is defined as

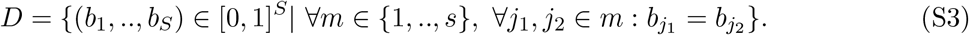

##### Proposition 1.

Assume that the trade-off relation is strictly concave (i.e., *γ* > 1). Then there is a unique colony phenotype 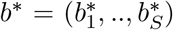 that represents the point of local maximum (on the set *D*) of the fitness function defined by Equation S1. This phenotype can be found in the following way:

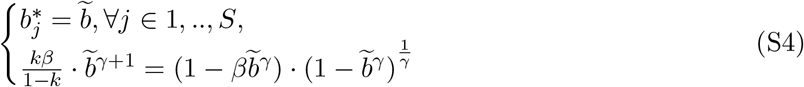

i.e., we conclude that for *G* → ∞, a population of small colonies evolves to a state at which all cells have the same unspecialized phenotype 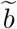 . The proof of Proposition 1 is presented in Section 5.2.4.

In general, Equation S4 can be solved numerically. However, we can get some analytical progress for the number of special cases of the model.

**The case of** *β* = 1. With *β* = 1, the equilibrium cell phenotype, 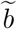, and the equilibrium population size, *N* ^*\**^, can be found explicitly:

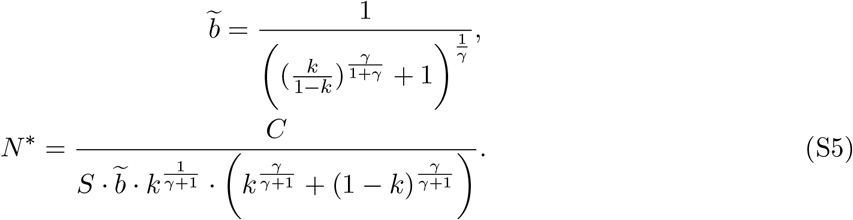

**Figure S9:**
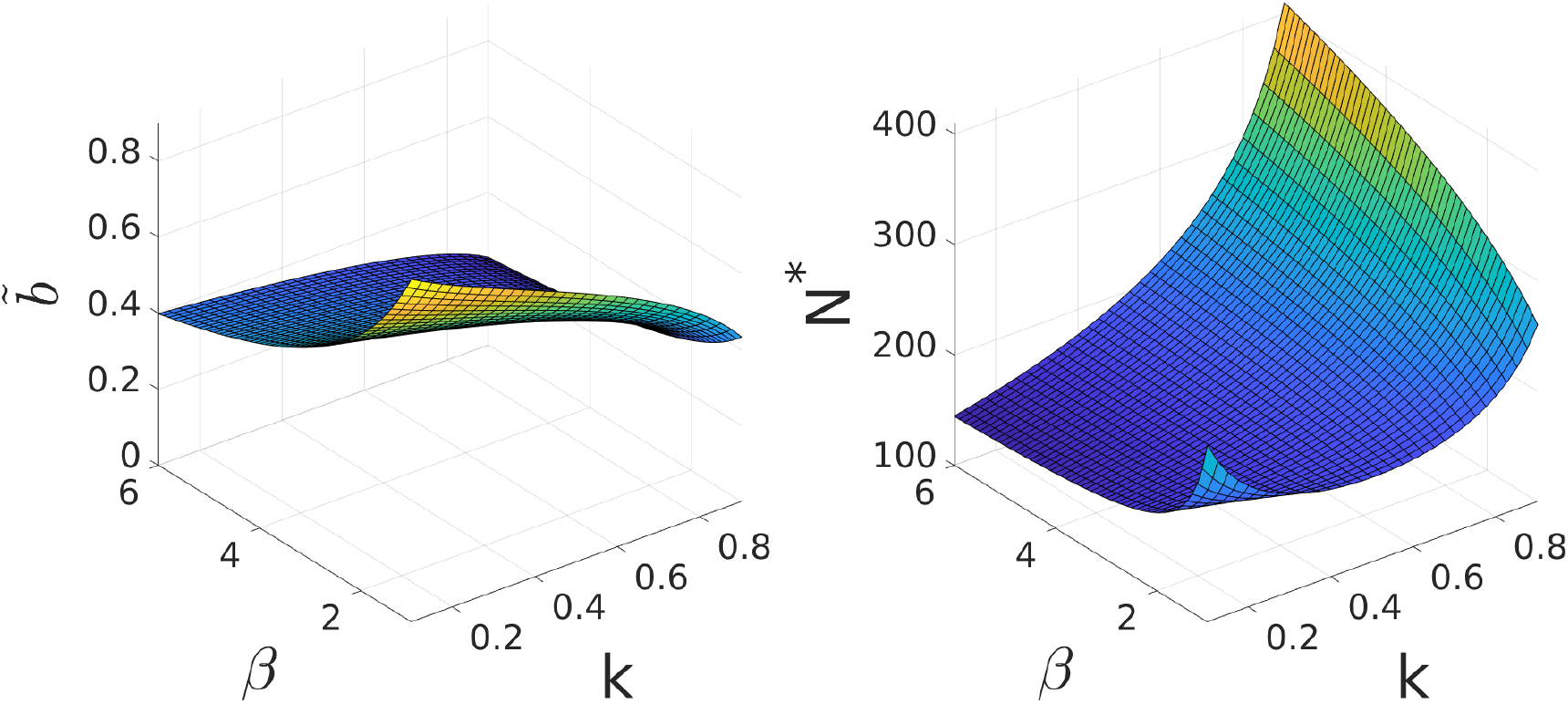
Analytical results for small colonies, *G* → ∞ and *γ* = 2. Effects of the relative costs of fecundity *k* and the strength of competition *β* on the equilibrium cell phenotype 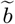 (the left panel) and the equilibrium population size *N* ^*\**^ (the right panel). Baseline parameter: *C* = 1000, *S* = 8.

Effects of the relative cost of fecundity *k* and the shape of the trade-off relation *γ* on the equilibrium cell phenotype 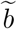 and the equilibrium population size *N* ^*\**^ are shown in Figure S8. Specifically, increasing *γ* increases 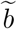 and decreases *N* ^*\**^. Increasing *k* leads to a decrease in 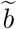 . The equilibrium population size *N* ^*\**^ exhibits a U-shaped dependence on *k*.

**The case of** *γ* = 2. Applying the Cardano formula, we get the following results:

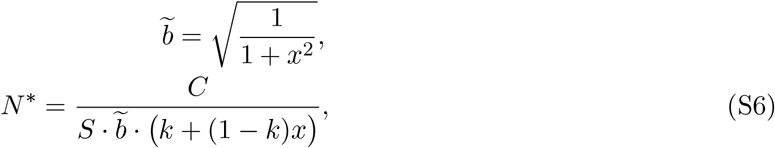

where *x* is a unique non-negative real number, such that:

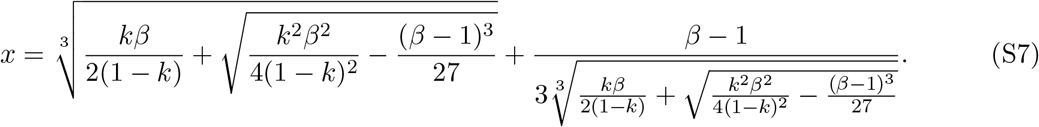

Effects of the relative cost of fecundity *k* and the strength of competition *β* on the equilibrium cell phenotype 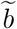 and the equilibrium population size *N* ^*\**^ are shown in Figure S9. Increasing *β* or increasing *k* decrease 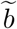 . Increasing *β*: (1) decreases *N* ^*\**^, if *k* is small; (2) has no significant effect on *N* ^*\**^, if *k* is intermediate; and (3) increases *N* ^*\**^, if *k* is large. The effect of *k* on *N* ^*\**^ depends on *β*. For small *β* the equilibrium population size *N*^*\**^ exhibits a U-shaped dependence on *k*. For intermediate or large *β* increasing *k* leads to an increase in *N*^*\**^.

##### General results

The results reported so far are summarized in Tables S1 and S2. Equation S4 was solved numerically to obtain the equilibrium cell phenotype 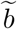 and the equilibrium population size *N* ^*\**^ for the cases with 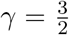, *β* = 3 and 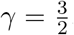, *β* = 6 which are not covered by our analytical analysis.

**Table S1:** Analytical results for small colonies and *G* → ∞: the equilibrium cell phenotype 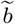.

|  |  | k=0.1 | k=0.2 | k=0.3 | k=0.4 | k=0.5 | k=0.6 | k=0.7 | k=0.8 | k=0.9 |
| --- | --- | --- | --- | --- | --- | --- | --- | --- | --- | --- |
| $\beta = 1$ | $\gamma = 2$ | 0.901 | 0.846 | 0.798 | 0.753 | 0.707 | 0.658 | 0.602 | 0.533 | 0.433 |
| | $\gamma = \frac{3}{2}$ | 0.854 | 0.786 | 0.731 | 0.68 | 0.63 | 0.578 | 0.521 | 0.451 | 0.355 |
| $\beta = 3$ | $\gamma = 2$ | 0.557 | 0.536 | 0.515 | 0.492 | 0.467 | 0.439 | 0.405 | 0.361 | 0.296 |
| | $\gamma = \frac{3}{2}$ | 0.461 | 0.441 | 0.42 | 0.399 | 0.376 | 0.35 | 0.32 | 0.28 | 0.223 |
| $\beta = 6$ | $\gamma = 2$ | 0.399 | 0.388 | 0.377 | 0.367 | 0.349 | 0.331 | 0.308 | 0.278 | 0.231 |
| | $\gamma = \frac{3}{2}$ | 0.296 | 0.288 | 0.279 | 0.269 | 0.257 | 0.242 | 0.225 | 0.201 | 0.163 |

**Table S2:** Analytical results for small colonies and *G* → ∞: the equilibrium population size *N* ^*\**^ (*C* =1000).

|  |  | k=0.1 | k=0.2 | k=0.3 | k=0.4 | k=0.5 | k=0.6 | k=0.7 | k=0.8 | k=0.9 |
| --- | --- | --- | --- | --- | --- | --- | --- | --- | --- | --- |
| $\beta = 1$ | $\gamma = 2$ | 260 | 210 | 189 | 180 | 177 | 180 | 189 | 210 | 260 |
| | $\gamma = \frac{3}{2}$ | 155 | 121 | 107 | 101 | 99 | 101 | 107 | 121 | 155 |
| $\beta = 3$ | $\gamma = 2$ | 156 | 160 | 166 | 174 | 185 | 201 | 224 | 263 | 345 |
| | $\gamma = \frac{3}{2}$ | 84 | 87 | 90 | 96 | 103 | 113 | 128 | 155 | 213 |
| $\beta = 6$ | $\gamma = 2$ | 144 | 154 | 164 | 176 | 194 | 217 | 250 | 302 | 409 |
| | $\gamma = \frac{3}{2}$ | 75 | 81 | 88 | 96 | 107 | 122 | 143 | 179 | 258 |

#### 5.2.3 Fitness landscape analysis for a population of large colonies

Here we assume that in large colonies the trade-off function describing the relationship between activity *a* and fecundity *b* is convex with *γ <* 1.

##### Proposition 2

Assume that the trade-off function is strictly convex (i.e.,*γ <* 1). Then there is no more than one unspecialized cell prototype at a point of local maximum (on *D*) of the fitness function defined by Equation S1.

A proof of Proposition 2 is presented in Section 5.2.4. Let us introduce the following notation. We say that a vector (*n*_1_, *n*_2_, *n*_3_), where *n*_1_, *n*_2_, *n*_3_ ∈ {0, 1, .., *s*} and *n*_1_ +*n*_2_ +*n*_3_ = *s*, is a phenotypic configuration of a colony if and only if *n*_1_ is the number of reproductive cell prototypes; *n*_3_ is the number of somatic cell prototypes; and *n*_2_ is the number of unspecialized cell prototypes. We say that a configuration (*n*_1_, *n*_2_, *n*_3_) is full-specialized, if *n*_2_ = 0. Proposition 2 states that in the equilibrium phenotypic configuration *n*_2_ ∈ {0, 1}. For further analytical progress, consider a special case of the model.

##### Special case

Consider a colony of *S* = 32 cells divided equally between *s* = 4 prototypes. Let 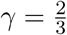, *β* ∈ {1, 3, 6} and *k* ∈ {0.1, 0.2, 0.3, 0.4, 0.5, 0.6, 0.7, 0.8, 0.9}. According to Proposition 2 one concludes that the equilibrium colony phenotype can be characterized by one of the following 7 configurations: {(3, 1, 0), (3, 0, 1), (2, 1, 1), (2, 0, 2), (1, 1, 2), (1, 0, 3), (0, 1, 3)}.

Numerical calculations (see Figure S11) show that an equilibrium phenotypic configuration is full-specialized except cases with *β* = 6 and *k* ∈ {0.8, 0.9} (where colonies are characterized by the configuration with the unique unspecialized prototype, (0,1,3)). Therefore, in general, an equilibrium phenotypic configuration can be found among full-specialized configurations. Calculating fitness of full-specialized configurations directly, we get:

- The configuration (3,0,1) has the highest fitness among all full-specialized configurations if and only if 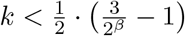.
- The configuration (2,0,2) has the highest fitness among all full-specialized configurations if and only if 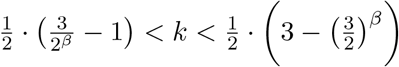.
- The configuration (1,0,3) has the highest fitness among all full-specialized configurations if and only if 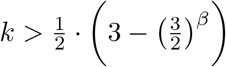.

**Figure S10:**
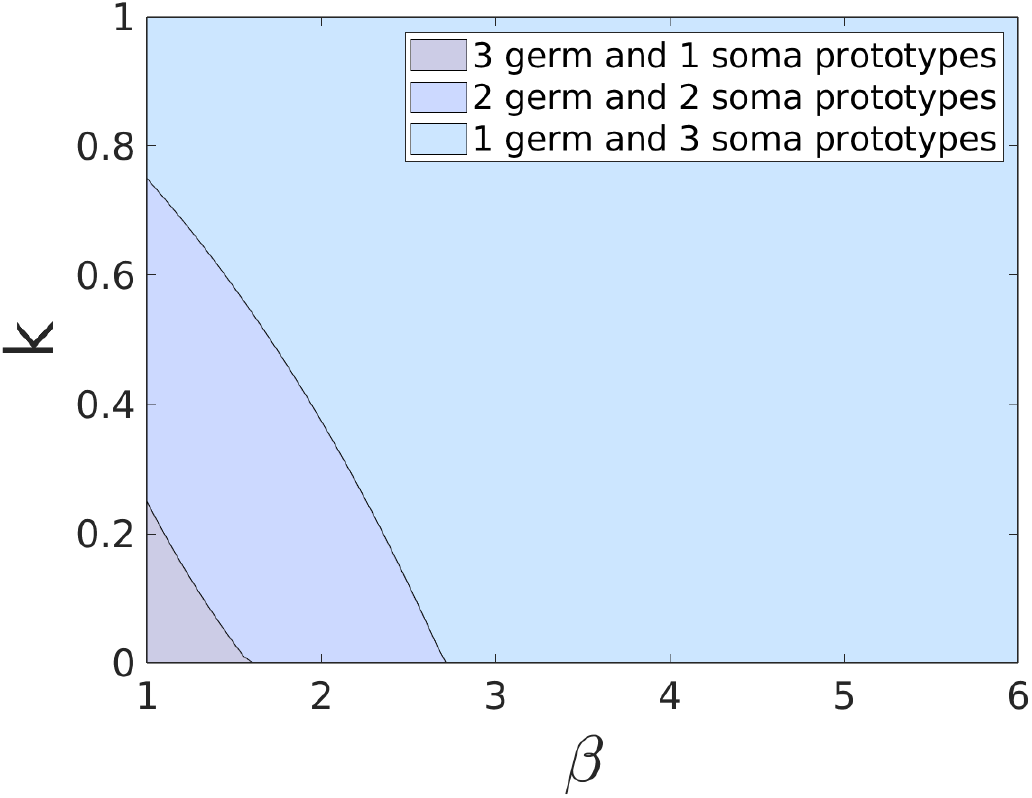
Analytical results for large colonies and *G* → ∞. Effects of the relative costs of fecundity *k* and the strength of competition *β* on a phenotypic configuration with the highest fitness among all full-specialized configurations. Baseline parameter: *C* = 1000, *S* = 32, 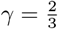, *s* = 4, there are 8 cells of each prototype in a colony.

These results are illustrated in Figure S10. One concludes that: (1) the configuration (3,0,1) has the highest fitness among all full-specialized configurations, if *k* and *β* are small; (2) (1,0,3) has the highest fitness, if *k* is large or *β* is intermediate or large; and (3) for all other cases (2,0,2) has the highest fitness.

#### 5.2.4 Proofs

**Proof of Proposition 1**. First, let us show that all coordinates of each point of local maximum of *W* on *D* are equal. To do this consider a reproductive strategy of a new colony in a resident population *b*′ ∈ *D*, such that two prototypes, *m*_1_ and *m*_2_ have different fecundities: 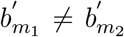 . Consider another reproductive strategy, *b*″ ∈ *D* such that 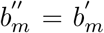 for all prototypes except *m*_1_ and *m*_2_, 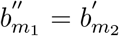 and 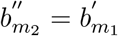 . Consider also all reproductive strategies *b*^*λ*^ = *λb*′ + (1 − *λ*)*b*″, where *λ* ∈ (0, 1). Since *D* is convex, all these strategies are available to the colony. Note that ∀*λ* ∈ (0, 1) : *B*(*b*′) = *B*(*b*″) = *B*(*b*^*λ*^); and ∀*λ* ∈ (0, 1) : *A*(*b*′) = *A*(*b*″) and *A*(*b*^*λ*^) > *A*(*b*′) (due to the strict concavity of the trade-off function). Let’s calculate the difference between fitness obtained by a strategy *b*^*λ*^, *λ* ∈ (0, 1) and that obtained by the strategy *b*′ :

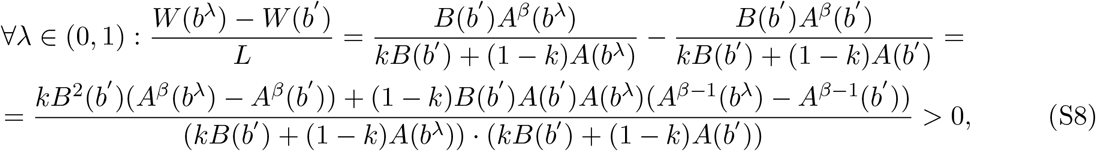

where 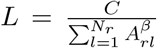 . It implies that for the strategy *b*′ and each *δ* > 0 we can find a strategy *b*^*λ*^, *λ* ∈ (0, 1) that belongs to an open ball *U*_*δ*_(*b*′) ∩ *D* and yields a higher fitness than the strategy*b*′.

Second, we consider the behavior of the fitness function on the set of all available strategies (*b*, .., *b*), such that all cells within the colony have the same fecundity *b*:

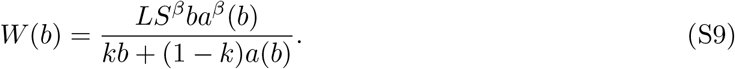

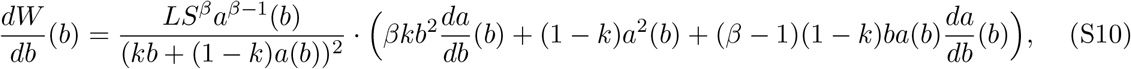

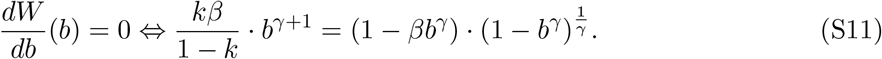

The last equation has the unique solution. To show this, denote 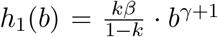 and 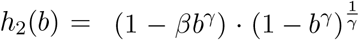 . The function *h*_1_(*b*) is continuous, strictly increasing and non-negative on the interval (0, 1). The function *h*_2_(*b*) is continuous and strictly decreasing on the interval (0, *θ*); and continuous and strictly increasing on the interval (*θ*, 1), where 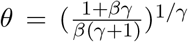 . Moreover, *h*_1_(0) = 0, *h*_2_(0) = 1, *h*_1_(*θ*) > 0, *h*_2_(*θ*) *<* 0 and *h*_2_(1) = 0. Therefore, one concludes that there exists the unique solution 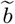 of Equation S11 on the interval (0, 1); and this solution belongs to the interval (0, *θ*).

Finally, since *W* (0, .., 0) = *W* (1, …, 1) and *W* (*b*) > 0 ∀*b* ∈ *D*\{(0, .., 0), (1, …, 1)}, the strategy 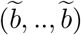 yields a higher fitness to the colony than other available strategies with identical fecundities of all cells. Since the function *W* is continuous on the compact set *D*, 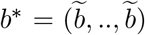 is the unique point of local and global maximum of the function *W* on *D*. **The proposition is proved**.

**Proof of Proposition 2**. Let *b*′ ∈ *D* be a reproductive strategy of a new colony in a resident population such that two prototypes, *m*_1_ and *m*_2_ are unspecialized i.e., 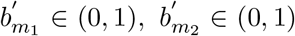 . Our goal is to show that the point *b*′ is not a point of local maximum (on *D*) of the fitness function *W* . Indeed, for each 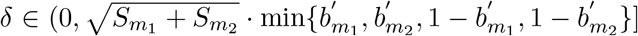 we find a reproductive strategy from *D* ∩ *U*_*δ*_(*b*′) that yields a higher fitness than the strategy *b*′ . To do this, consider two reproductive strategies, *b*″, *b*‴ ∈ *D* such that 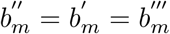 for all prototypes except *m*_1_ and *m*_2_, 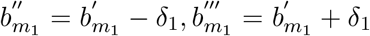 and 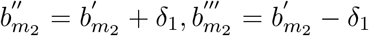, where 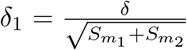 . Note that *b*″, *b*‴ ∈ *D* ∩ *U* (*b*′). Moreover, *B*(*b*′) = *B*(*b*″) = *B*(*b*‴) and 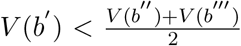 due to the strict convexity of the trade-off function. It implies that *V* (*b*′) *< V* (*b*″) or *V* (*b*′) *< V* (*b*‴). Without loss of generality, we postulate that *V* (*b*′) *< V* (*b*″). Repeating the arguments from the proof of Proposition 1, we conclude that *W* (*b*″) > *W* (*b*′). **The proposition is proved**.

#### 5.2.5 Additional numerical results for the case of *G* → ∞

**Figure S11:**
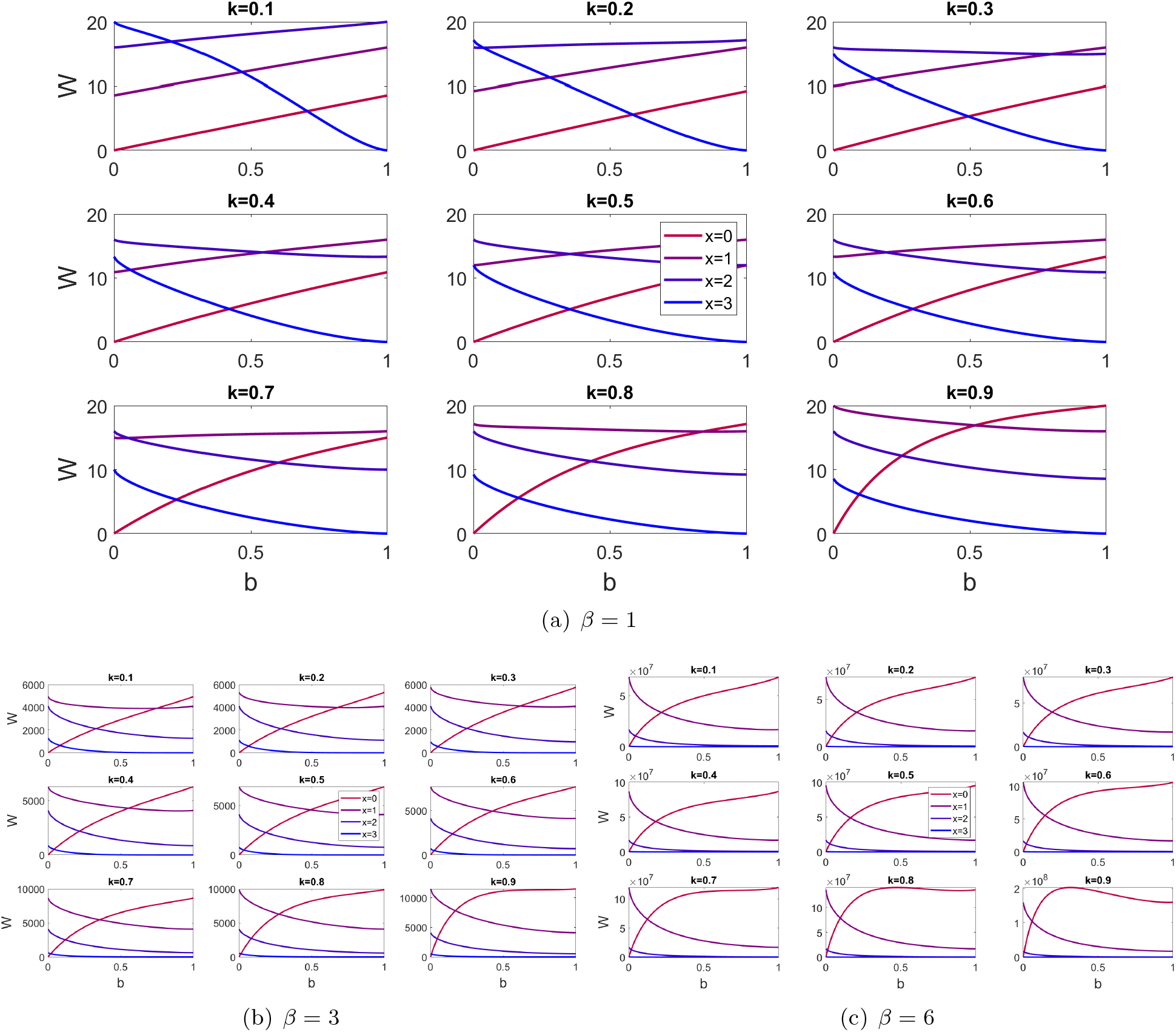
Fitness landscape for a colony with *x* reproductive, 3 − *x* somatic (*x* ∈ {0, 1, 2, 3}), and one variable (with fecundity *b*) prototypes. *S* = 32, 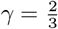, *s* = 4, *L* = 1, there are 8 cells of each prototype in a colony.

## References

Amado, A., Batista, C., and Campos, P. (2017). A theoretical approach to the size-complexity rule. Evolution, 72-1, 18–29.

Amado, A., Batista, C., and Campos, P. (2018). A mechanistic model for the evolution of multi-cellularity. Physica A: Statistical Mechanics and its Applications, 492, 1543–1554.

Berman-Frank, I., Lundgren, P., and Falkowski, P. (2003). Nitrogen fixation and photosynthetic oxygen evolution in cyanobacteria. Research in microbiology, 154(3), 157–164.

Beshers, S. N. and Fewell, J. H. (2001). Models of division of labor in social insects. Annual Reviews in Entomology, 46, 413–440.

Bonner, J. (2003). Evolution of development in the cellular slime molds. Evolution and Devlopment, 5, 305–313.

Bonner, J. T. (2004). Perspective: The size-complexity rule. Evolution, 58, 1883–1890.

Brumley, D., Polin, M., Pedley, T., and Goldstein, R. (2015). Metachronal waves in the flagellar beating of volvox and their hydrodynamic origin. Journal of the Royal Society Interface, 12(108), 20141358.

Carroll, S. (2001). Chance and necessity: the evolution of morphological complexity and diversity. Nature, 409(6823), 1102–1109.

Cooper, G. and West, S. (2018). Division of labour and the evolution of extreme specialization. Nature ecology & evolution, 2(7), 1161–1167.

de Jong, J. (1990). Quantitative genetics of reaction norms. Journal of Evolutionary Biology, 3, 447–468.

de Oliveira, V., Mendes, B., Roque, M., and Campos, P. (2020). Extinction-colonization dynamics upon a survival-dispersal trade-off. Ecological Complexity, 43, 100856.

Duarte, A., Weissing, F., Pen, I., and Keller, L. (2011). An evolutionary perspective on selforganized division of labor in social insects. Annual review of ecology, evolution, and systematics, 42, 91–110.

Durkheim, E. (2014). The division of labor in society. Simon and Schuster.

Egas, M., Dieckmann, U., and Sabelis, M. (2004). Evolution restricts the coexistence of specialists and generalists: the role of trade-off structure. American Naturalist, 163(4), 518–531.

Fay, P. (1992). Oxygen relations of nitrogen fixation in cyanobacteria. Microbiology and Molecular Biology Reviews, 56(2), 340–373.

Flores, E. and Herrero, A. (2010). Compartmentalized function through cell differentiation in filamentous cyanobacteria. Nature Reviews Microbiology, 8(1), 39–50.

Folse, H. J. and Roughgarden, J. (2010). What is an individual organism? A multilevel selection perspective. Quarterly Reviews of Biology, 85, 446–472.

Gavrilets, S. (2010). Rapid transition towards the division of labor via evolution of developmental plasticity. Plos Computational Biology, 6.

Gavrilets, S. and Scheiner, S. (1993). The genetics of phenotypic plasticity. v. evolution of reaction norm shape. Journal of Evolutionary Biology, 6, 31–48.

Goebl, M. and Petes, T. (1986). Most of the yeast genomic sequences are not essential for cell growth and division. Cell, 46(7), 983–992.

Goldsby, H., Knoester, D., Ofria, C., and Kerr, B. (2014). The evolutionary origin of somatic cells under the dirty work hypothesis. PLoS Biology, 12(5), e1001858.

Goldstein, R. (2015). Green algae as model organisms for biological fluid dynamics. Annual Review of Fluid Mechanics, 47, 343–375.

Grosberg, R. K. and Strathmann, R. R. (2007). The evolution of multicellularity: A minor major transition? Annual Review of Ecology and Systematics, 38, 621–654.

Herron, M., Ghimire, S., Vinikoor, C., and Michod, R. (2014). Fitness trade-offs and developmental constraints in the evolution of soma: An experimental study in a volvocine algae. Ecology and Evolutionary Biology, 16, 203–221.

Herron, M. D. and Michod, R. E. (2007). Evolution of complexity in the volvocine algae: transition in individuality through Darwin’s eye. Evolution, 62, 436–451.

Ispolatov, I., Ackermann, M., and Doebeli, M. (2012). Division of labour and the evolution of multicellularity. Proceedings of the Royal Society London B, 279, 1768–1776.

Kaufman, L. and Rousseeuw, P. (1990). Finding Groups in Data: An Introduction to Cluster Analysis. John Wiley & Sons, USA.

Kirk, D. (2001). Germ-soma differentiation in Volvox. Developmental Biology, 238(2), 213–223.

Kirk, D. (2003). Seeking the ultimate and proximate causes of Volvox multicellularity and cellular differentiation. Integrative and Comparative Biology, 43(2), 247–253.

Kirk, D. (2005). A twelve-step program for evolving multicellularity and a division of labor. Bioessays, 27(3), 299–310.

Konrad, K. A. (2009). Strategy and dynamics in contests. Oxford University Press, Oxford, United Kingdom.

Leslie, M., Shelton, D., and Michod, R. (2017). Generation time and fitness tradeoffs during the evolution of multicellularity. Journal of Theoretical Biology, 430, 92–102.

Libby, E. and Ratcliff, W. (2014). Ratcheting the evolution of multicellularity. Science, 346(6208), 426–427.

McShea, D. (2000). Functional complexity in organisms: Parts as proxies. Biology and Philosophy, 15, 641–668.

Michod, R. (1996). Cooperation and conflict in the evolution of individuality. Proceedings of the Royal Society London B, 263(1383), 813–822.

Michod, R. (1997). Evolution of the individual. American Naturalist, 150(Suppl. S), S5–S21.

Michod, R. (2005). On the transfer of fitness from the cell to the multicellular organism. Biology and Philosophy, 20(5), 967–987.

Michod, R., Viossat, Y., Solari, C., Hurand, M., and Nedelcu, A. (2006). Life-history evolution and the origin of multicellularity. Journal of Theoretical Biology, 239(2), 257–272.

Michod, R. E. (2006). The group covariance effect and fitness trade-offs during evolutionary transitions in individuality. Proceedings of the National Academy of Sciences USA, 103(24), 9113–9117.

Michod, R. E. (2007). Evolution of individuality during the transition from unicellular to multicellular life. Proceedings of the National Academy of Sciences USA, 104(Suppl. 1), 8613–8618.

Nedelcu, A. (2009). Environmentally induced responses co-opted for reproductive altruism. Biology Letters, 5, 805–808.

Nedelcu, A. and Michod, R. (2006). The evolutionary origin of an altruistic gene. Molecular Biology and Evolution, 23(8), 1460–1464.

Oliver, S., Van der Aart, Q., Agostoni-Carbone, M., Aigle, M., Alberghina, L., Alexandraki, D., Antoine, G., Anwar, R., Ballesta, J., Benit, P., et al. (1992). The complete dna sequence of yeast chromosome iii. Nature, 357(6373), 38–46.

Panter-Brick, C. (2002). Sexual division of labor: energetic and evolutionary scenarios. American Journal of Human Biology, 14(5), 627–640.

Pichugin, Y. and Traulsen, A. (2020). Evolution of multicellular life cycles under costly fragmentation. PLoS computational biology, 16(11), e1008406.

Pichugin, Y., Peña, J., Rainey, P., and Traulsen, A. (2017). Fragmentation modes and the evolution of life cycles. Plos Computational Biology, 13(11).

Pichugin, Y., Park, H., and Traulsen, A. (2019). Evolution of simple multicellular life cycles in dynamic environments. Journal of the Royal Society Interface, 16(154), 20190054.

Reuffler, C. and Wagner, G. (2012). Evolution of functional specialization and division of labor. Proceedings of the National Academy of Sciences USA, 109(6), E326–E335.

Robinson, G. E. (1992). Regulation of division of labor in insect societies. Annual Review of Entomology, 37, 637–665.

Rodriguez-Clare, A. (1996). The division of labor and economic development. Journal of Development Economics, 49, 3–32.

Rossetti, V., Schirrmeister, B. E., Bernasconi, M. V., and Bagheri, H. C. (2010). The evolutionary path to terminal differentiation and division of labor in cyanobacteria. Journal of Theoretical Biology, 262, 23–34.

Rusch, H. and Gavrilets, S. (2017). The logic of animal intergroup conflict: a review. Journal of Economic Behavior & Organization.

Shelton, D., Desnitskiy, A., and Michod, R. (2012). Distributions of reproductive and somatic cell numbers in diverse volvox (chlorophyta) species. Evolutionary ecology research, 14, 707.

Smith, A. (2008). An Inquiry into the Nature and Causes of the Wealth of Nations: A Selected Edition Adam Smith (Author). Kathryn Sutherland (Editor), Oxford Paperbacks, Oxford.

Smith, J. and Szathmary, E. (1995). The Major Transitions in Evolution. Freeman, San Francisco.

Staps, M., van Gestel, J., and Tarnita, C. (2019). Emergence of diverse life cycles and life histories at the origin of multicellularity. Nature ecology & evolution, 3(8), 1197–1205.

Stigler, G. (1951). The division of labor is limited by the extent of the market. Journal of political economy, 59(3), 185–193.

Thatcher, J., Shaw, J., and Dickinson, W. (1998). Marginal fitness contributions of nonessential genes in yeast. Proceedings of the National Academy of Sciences, 95(1), 253–257.

Tullock, G. (1980). Efficient rent seeking. In J.M. Buchanan, R.D. Tollison, and G. Tullock, editors, Toward a theory of the rent-seeking society, page 97–112. Texas A & M University, College Station.

Tverskoi, D., Makarenkov, V., and Aleskerov, F. (2018). Modeling functional specialization of a cell colony under different fecundity and viability rates and resource constraint. Plos One, 13(8).

West, S., Fisher, R., Gardner, A., and Kiers, E. (2015). Major evolutionary transitions in individuality. Proceedings of the National Academy of Sciences, 112(33), 10112–10119.

Willensdorfer, M. (2009). On the evolution of differentiated multicellularity. Evolution, 63(2), 306–323.

Wilson, E. O. (2008). One giant leap: how insects achieved altruism and colinial life. BioScience, 58, 17–25.

Yanni, D., Jacobeen, S., Márquez-Zacarías, P., Weitz, J., Ratcliff, W., and Yunker, P. (2020). Topological constraints in early multicellularity favor reproductive division of labor. Elife, 9, e54348.

